# Ancient *Borrelia* genomes document the evolutionary history of louse-borne relapsing fever

**DOI:** 10.1101/2024.07.18.603748

**Authors:** Pooja Swali, Thomas Booth, Cedric C.S. Tan, Jesse McCabe, Kyriaki Anastasiadou, Christopher Barrington, Matteo Borrini, Adelle Bricking, Jo Buckberry, Lindsey Büster, Rea Carlin, Alexandre Gilardet, Isabelle Glocke, Joel Irish, Monica Kelly, Megan King, Fiona Petchey, Jessica Peto, Marina Silva, Leo Speidel, Frankie Tait, Adelina Teoaca, Satu Valoriani, Mia Williams, Richard Madgwick, Graham Mullan, Linda Wilson, Kevin Cootes, Ian Armit, Maximiliano G. Gutierrez, Lucy van Dorp, Pontus Skoglund

## Abstract

Several disease-causing bacteria have transitioned from tick-borne to louse-borne transmission, a process associated with increased virulence and genome reduction. However, the historical time frame and speed of such evolutionary transitions have not been documented with ancient genomes. Here, we discover four ancient cases of *Borrelia recurrentis*, the causative agent of louse-borne relapsing fever, in Britain between ∼600 and 2,300 years ago, and sequence whole genomes up to 29-fold coverage. We estimate a recent divergence from the closest tick-borne ancestor, likely within the last ∼8,000 years. We reconstruct a chronology of gene losses and acquisitions using the pan-genome of related species, and show that almost all of the reductive evolution observed in *B. recurrentis* had occurred by ∼2,000 years ago, and was thus a rapid process after divergence. Our observations provide a new understanding of the origins of *B. recurrentis* and document complex reductive evolution in a specialist vector-borne pathogen.

## Introduction

Several species of bacteria have undergone an evolutionary process of transitioning from tick-borne to louse-borne transmission, including the trench fever agent *Bartonella quintana*, the epidemic typhus agent *Rickettsia prowazekii*, and the agent of louse-borne relapsing fever (LBRF) *Borrelia recurrentis*. All species show a pattern of higher virulence in the louse-borne agent compared to their respective closest tick-borne relatives, and all show a striking evolutionary pattern of genome reduction^1^, possibly facilitated by specialisation to the louse vector^1–3^. However, the evolutionary time frame and genomic basis of the transition from tick-borne to louse-borne transmission, and the drivers of increased virulence, remain largely unknown.

Relapsing fever, named after the recurring fevers it induces, is caused by several species of *Borrelia.* These are mostly spread by soft-bodied ticks, with the exception of *Borrelia miyamotoi* which is spread by hard-bodied ticks (tick-borne relapsing fever; TBRF)^4^ and *B. recurrentis,* which is transmitted from human to human by the bite of an infected human body louse, *Pediculus humanus humanus*^5^; the latter is not known to have an animal reservoir. The bacteria establishes infection when the haemocoel of the infected vector is able to penetrate the mucosa membrane or skin barrier^5^. In contrast to LBRF, TBRF is zoonotic, with multiple animal reservoir hosts, and can be found worldwide. For example, the closest related bacterial species to *B. recurrentis*, *Borrelia duttonii*, was previously found in animal reservoirs such as pigs and chickens and is today mostly found in East Africa^6,7^.

The present-day *B. recurrentis* genome has an unusual genome structure, comprising a ∼930 kb linear chromosome and seven linear plasmids ranging from 6-124 kb in length^8^. Though the chromosome is fairly conserved over the *Borrelia* genus, the plasmids have potential to undergo extensive rearrangements^3^. *B. recurrentis* has been estimated to have lost approximately a fifth of its genome relative to its sister species *B. duttonii*, with prominent gene loss occurring on plasmids^3^. It has been suggested that genome loss in the other louse-borne taxa in *Rickettsia* and *Bartonella* was primarily via elimination of inactivated genes^1,3^, which has also been suggested for *B. recurrentis*. The exact genes involved in the mechanism for vector specification (louse or tick), and the evolutionary processes underlying genome degradation in *Borrelia*, remain unclear.

Major uncertainties surround the past and present epidemiology of *B. recurrentis* and hence the timeline over which genome reduction and vector/host specialisation occurred. Throughout European history, there have been numerous references to episodes of “epidemic fever”, and “fever lasting six or seven days, with multiple relapses”^9,10^; the earliest descriptions date back to ancient Greece in the 5th century BCE^10^. It has been hypothesised that LBRF was the agent of the Yellow Plague which affected Europe in 550 CE, the episodic fevers which became known as sweating sickness in northwestern Europe between 1,485-1,551 CE^11,12^, as well as fevers that accompanied famines in Ireland through the 17th and 18th centuries CE. However, the specific agents of these historical outbreaks have not been confirmed. LBRF posed major challenges to public health during World War I and World War II, before mostly disappearing from Europe at the end of the 20th century CE^13^. Today, LBRF remains a major cause of morbidity and mortality in Ethiopia (where it is endemic), Somalia and Sudan^13,14^. While some now consider LBRF as a neglected tropical disease (NTD), it is clear that LBRF shows the potential to re-emerge during times of overcrowding, poor access to sanitation and hygiene, and during times of conflict and disaster^3,15^.

*B. recurrentis* is a challenging species to grow in culture, so limited genomic data is available from present-day infections^8,16^. As such, an archaeogenetic approach represents one of the most promising tools for characterising the pathogen’s wider diversity and evolution. Previously, a ∼550-year-old (1,430–1,465 cal. CE, 95% confidence) 6.4-fold *B. recurrentis* genome was recovered from a tooth taken from a human skeleton buried in medieval Oslo, Norway (OSL9)^17^. However, we lack an understanding of the deeper genomic evolution of *B. recurrentis*, or its prevalence in Europe across time and space. Here we provide four new ancient *B. recurrentis* genomes from Britain spanning 2,300-600 years ago, from the Iron Age to the later medieval period. Leveraging these observations, we confirm the contribution of *B. recurrentis* to disease in European history and document its complex evolutionary behaviour during the transition to louse-borne transmission.

## Results

### Detection and authentication of four ancient *B. recurrentis* genomes

We used an ancient DNA approach, including single-stranded DNA library preparation^18^, which optimises retrieval of short fragments and allows postmortem cytosine deamination-derived errors to be removed *in-silico*, to generate whole *B. recurrentis* genomes from four human skeletons recovered from four archaeological sites in Britain (**Figure 1A**). We generated ∼0.8 - 8.5 billion read pairs, obtaining 0.8-29.4-fold coverage over the *B. recurrentis* A1 reference chromosome (**Table 1**). All libraries were sequenced to either more than 20-fold coverage or more than 40% clonality (the proportion of sequences identified as PCR duplicates, indicative of library saturation). These include two observations dating to the last ∼2,000 years: an 11.2-fold genome (C10416, Burial 240) from the ‘Arras-Culture’-associated Iron Age cemetery at Wetwang Slack, East Yorkshire, contextually dated to 2,300-2,100 years ago (300-100 BCE)^19^, and a 3.5-fold *B. recurrentis* genome from Fishmonger’s Swallet (C13361, mandible G10-1.4), a cave in southern Gloucestershire, UK. This latter individual has been directly radiocarbon dated to 2,185-2,033 years ago (162 cal BCE-10 CE; 95% confidence; 2063±28 BP, BRAMS-5059)^20^. We also generated a 0.8-fold *B. recurrentis* genome from the tooth of an unprovenanced skull (C11907, cranium CW29) from the collections of the Canterbury Archaeological Trust, which is likely to be from Canterbury or the surrounding region and dates to 2,000-700 years ago (1st-14th centuries CE). Finally, we generated a 29.4-fold *B. recurrentis* genome (C10976, Sk 435) from a rural cemetery site associated with a medieval chapel next to the village of Poulton, near Chester in Cheshire. The skeleton has been radiocarbon dated to 733-633 years ago (95% confidence, 1,290-1,390 cal CE, 646±14 BP, Wk 52986^21^. Further details of all individuals sampled in this study are available in the **Supplementary Information.**

**Figure 1.**
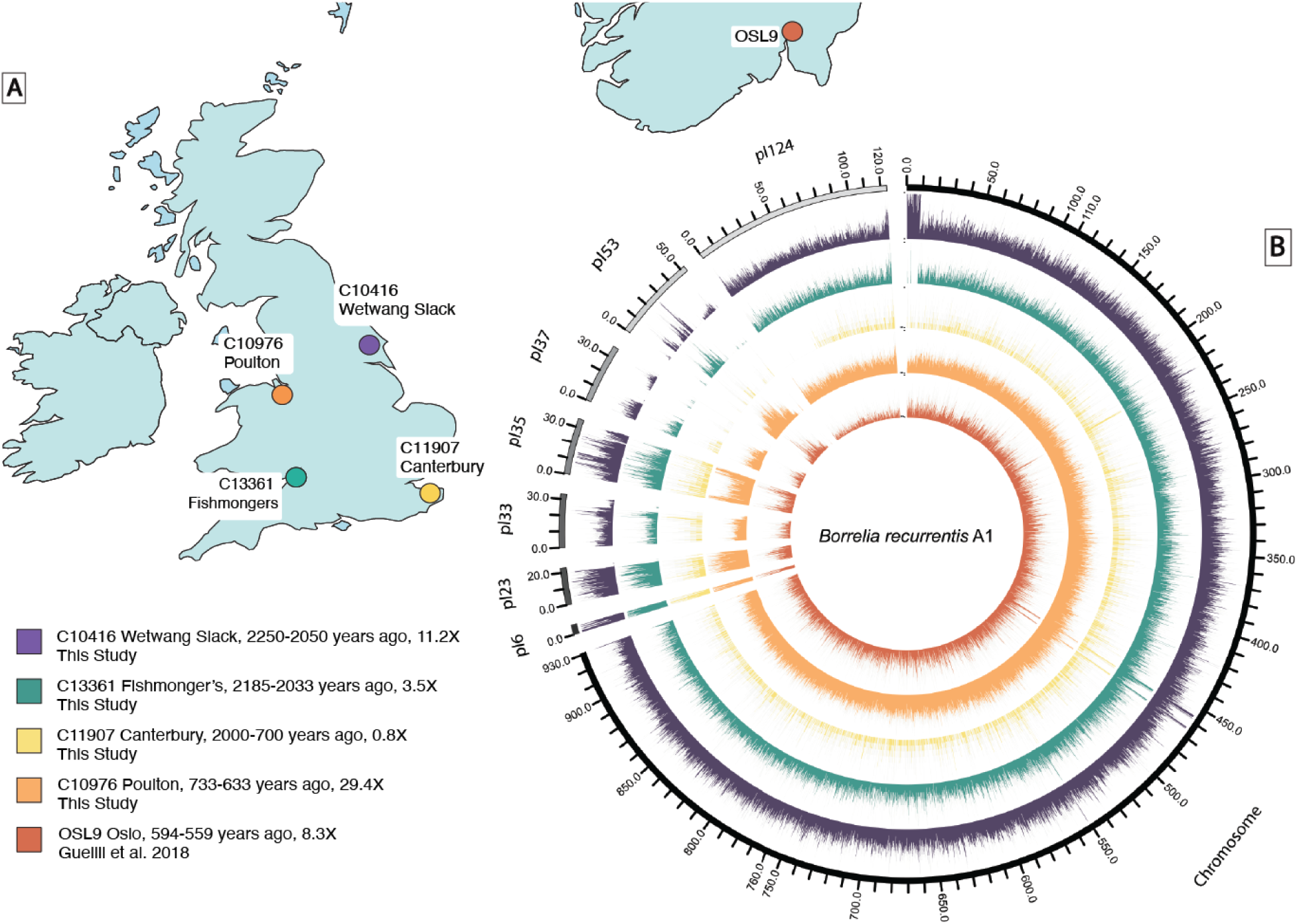
Overview of ancient genomes. A. Geographic location of the four ancient *B. recurrentis* genomes sequenced in this study together with OSL9 previously published by Guellil and colleagues^17^. **B.** Circos plot with the coverage of ancient genomes across the *B. recurrentis* chromosome and plasmids when aligned to the *B. recurrentis* A1 reference genome (GCF_000019705.1). A window size of 100bp for the chromosome and 10bp for the plasmids was used to provide the normalised coverage per window plotted. To allow for visualisation, the coverage for each genome was scaled by the maximum coverage per genome (C10416 Wetwang Slack, 70; C13361 Fishmonger’s, 20; C11907 Canterbury, 10; C10976 Poulton, 170; OSL9, 40).

**Table 1.**
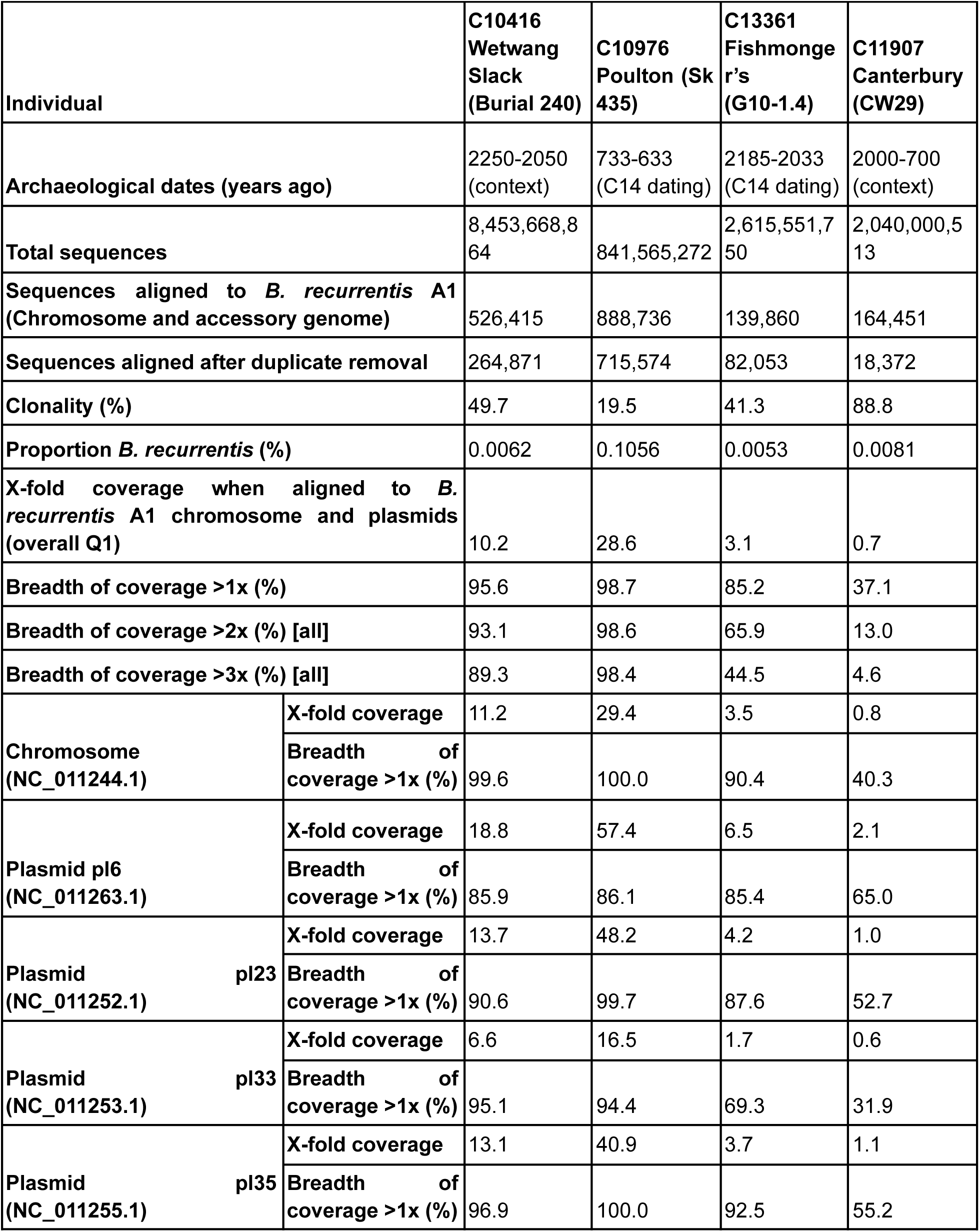
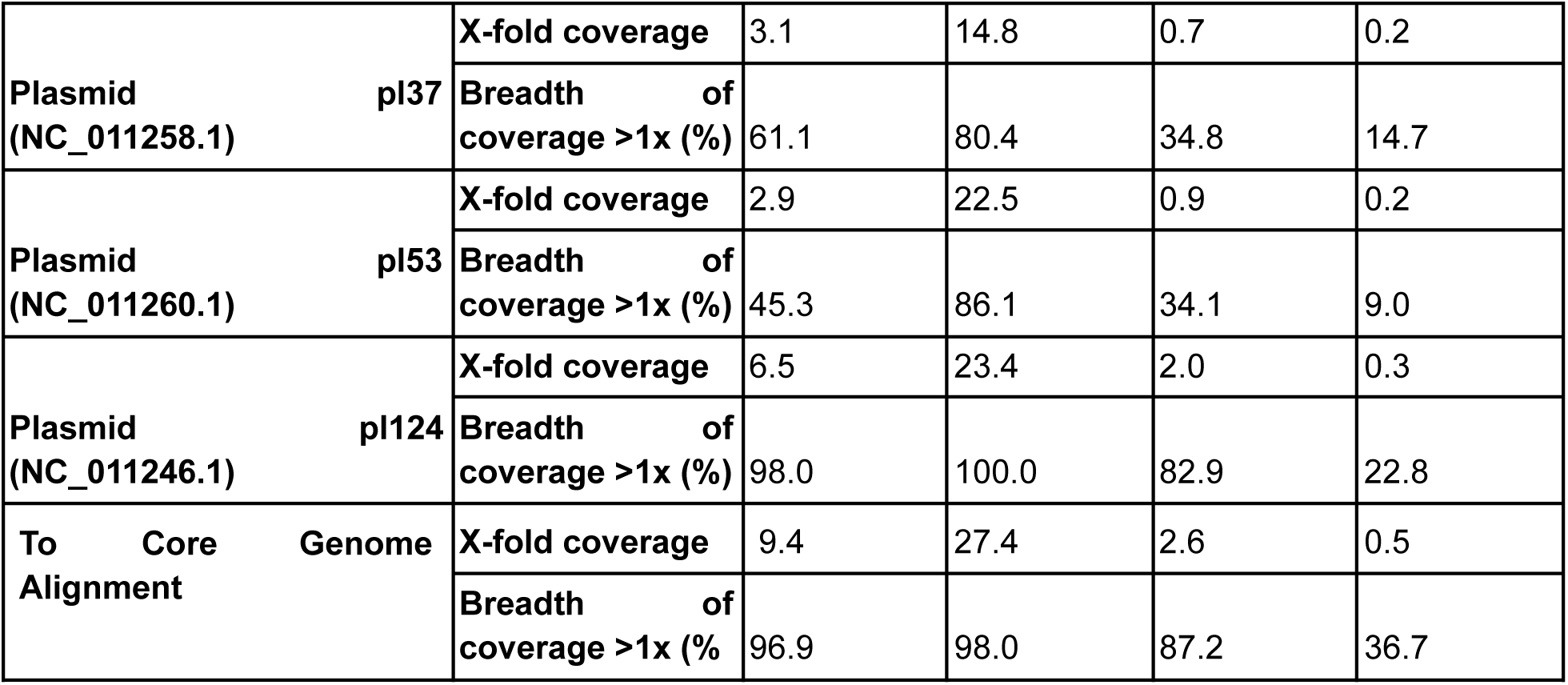
Sequencing metrics for the four individuals recovered in this study when mapped to the *B. recurrentis* A1 reference genome (chromosome and plasmids) requiring a mapping quality of MQ1.

Aligned sequences were confirmed to be authentic through assessment of evidence for cytosine deamination^22,23^, distribution of number of mismatches across sequences (edit distance), even coverage across the genome, and a unimodal fragment length distribution (**Figure 1B and Figure S1**)^24^. Additionally, all identified genomes were aligned to representatives of the more closely related species *B. duttonii* Ly and *Borrelia crocidurae* DOU reference strains, to confirm that all identified cases were genetically closer to the modern-day *B. recurrentis* A1 genome than other related *Borrelia* species (**Figure S2**). Mapping was also conducted to the wider *Borrelia* plasmid complement. The presence of the pl6 plasmid in these ancient British genomes, which is absent in *B. duttonii* Ly, further supports their taxonomic identification as *B. recurrentis*.

### Iron Age and medieval lineages of *B. recurrentis*

To evaluate the relatedness of our ancient strains to contemporary sampled strains, we initially reconstructed a *Borrelia* phylogeny including the closest *B. duttonii* representative (Ly). Consistent with the assessment of our samples being *B. recurrentis*, all ancient genomes form a monophyletic clade together with present-day *B. recurrentis* (**Figure S3 and Figure S4**). Our medieval genome, C10976 Poulton, is positioned in a subclade with the previously published medieval genome from Norway OSL9^17^. All Iron Age genomes from Britain fall basal to this clade. Among the Iron Age genomes, C13361 Fishmonger’s falls on a lineage basal to C10416 Wetwang Slack, despite being dated to a similar period. Although the Fishmonger’s genome is of lower coverage (2.6-fold when aligned to the core genome), the 100% bootstrap support for this phylogenetic placement suggests the possibility of synchronic sister lineages of the species existing in Britain ∼2,300-2,000 years ago. Additionally, we reconstructed a phylogeny on an alignment built using relaxed SNP filtering thresholds in order to include the lower coverage (0.8-fold) C11907 Canterbury genome (**Methods**) (**Figure S4**). We found this genome was closely related to Iron Age genomes C10416 Wetwang Slack and C13361 Fishmonger’s, yet the precise date of this individual is undetermined and ranges from 2,000-700 years ago. C11907 Canterbury was then excluded from further analysis due to its lower coverage (0.8-fold) and uncertain chronological date.

We next reconstructed a core gene alignment to assess the extent to which recombination and accessory (plasmid) gene content may influence our reconstructed relationships, by identifying a set of genes shared amongst the modern sampled diversity of *B. recurrentis*. To do so, we applied the pan-genome analysis tool Panaroo^25^ to all modern *B. recurrentis* (seven genomes)*, B. duttonii* (two assemblies) and *B. crocidurae* (two assemblies) genomes (**Table S1**). The inferred *Borrelia* pan-genome comprised a total of 3,035 genes corresponding to a length of 2,223,831 base pairs. We observed a high degree of conservation within the *B. recurrentis* species, supporting a limited intraspecies pan-genome despite high plasmid carriage. Of these genes, we identified 933 as being present in 99% of included strains, providing a core gene reference panel to which the ancient samples were aligned before phylogenetic reconstruction. Phylogenies constructed by mapping to the *B. recurrentis* A1 reference genome and core-genome alignment showed identical phylogenetic topologies with improved bootstrap support, despite the latter providing a higher amount of reference diversity (4,192 SNPs versus 4,200 SNPs). This suggests a limited impact of mapping bias or structural variation on our observed patterns of relatedness.

Given that recombination may violate the assumptions of tree-building representations of diversity, we formally tested for evidence of homologous recombination using ClonalFrameML^26^. We observed very limited evidence of homologous recombination, with only a small fraction of the genome (<0.1%) estimated to derive from such processes. Nonetheless, after pruning the alignment for the modest amount of recombination detected by ClonalFrameML, we recovered a topologically identical phylogeny (**Figure S5**).

### Chronology of the divergence from the tick-borne sister species

The timeline over which *B. recurrentis* diverged from its common ancestor with *B. duttonii*, and subsequently evolved vector and host specificity, is uncertain. Here, we use our ancient *Borrelia* time series to calibrate the joint genealogical history of modern and ancient *B. recurrentis* and its mutation rate. We first tested a hypothesis of clock-like evolution in our core-genome alignment. We formally assessed the temporal structure in our recovered phylogenies using *BactDating*^27^ to test for a significant correlation between genomic diversity and sampling time using date randomisation (**Methods**). We assessed temporality including and excluding C13361 Fishmonger’s, due to its lower coverage, and obtained an R² of 0.69 (p-value 0.00080) and 0.66 (p-value 0.0011) respectively. This result suggests a significant temporal signal across our dataset (**Table S2**).

We next implemented formal Bayesian tip-dating calibration via *BEAST2*^28^ to provide a probabilistic assessment of the divergence of sampled *B. recurrentis* from the closest sequenced relative *B. duttonii* Ly. This approach jointly estimates the rate of mutation over the non-recombining fraction of the core alignment (**Methods**). Evaluating a suite of possible clock and demographic models, we estimate a split time of *B. recurrentis* genomes over 5-fold in coverage in our dataset from *B. duttonii* Ly ranging from 2,215-5,630 years ago. When the lower coverage C13361 Fishmonger’s sample is included, we estimate the split time from *B. duttonii* Ly ranges between 2,313-7,654 years ago, overlapping with the initial estimate (**Figure 2A and Figure S6, Figure S7, Table S3**). The best supported model indicates a divergence estimate of 5,156 (95% credible interval 4,724-5,630) years ago, corresponding to a rate of evolution of 5.0 x 10^-8^ (4.6 x 10^-8^ - 5.5 x 10^-8^ substitutions per site per year; 95% HPD values) (**Figure 2A**). Estimates from this model suggest an emergence of the Iron Age clade between 2,326-2,410 years ago with the medieval clade, including C10976 Poulton and the previously recovered OSL9 genome, dating to within the last 700 years.

**Figure 2.**
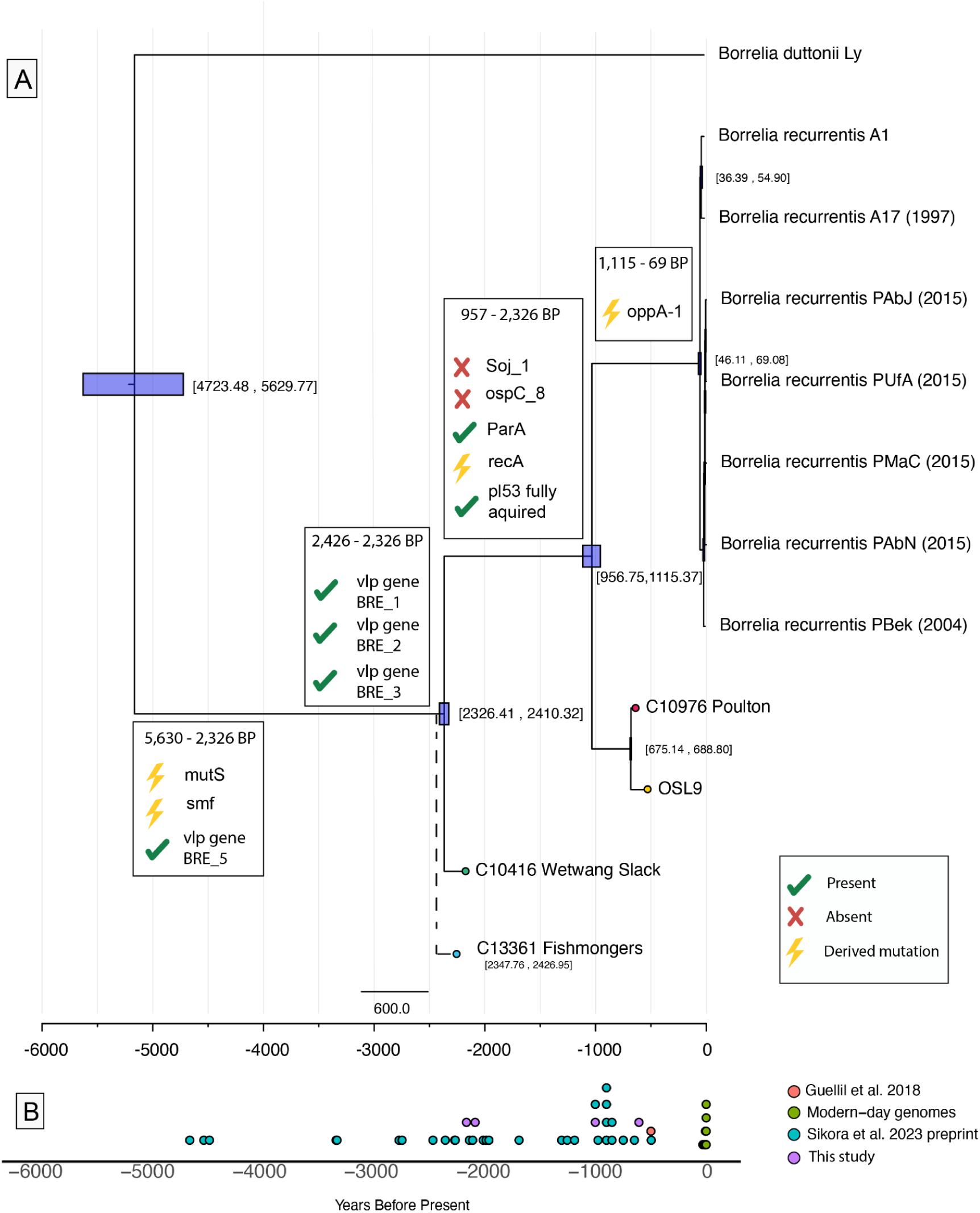
Temporal evolution of *Borrelia recurrentis*. A. Bayesian tip-calibrated maximum clade credibility time tree from Beast2, providing the best supported model following path-sampling. The 95% higher posterior density is indicated with purple boxes and within brackets. The placement of Fishmonger’s is indicative following a relaxed tip-calibration analysis. Ancient samples are highlighted by coloured tips. Key gene loss and gain events described in the text are highlighted at the relevant phylogenetic nodes. **B**. Timeline providing the estimated age of *B. recurrentis* observations recovered from ancient DNA reported in this and other studies.

Our inference would also suggest a very recent emergence (46-69 years ago) of all contemporarily sampled *B. recurrentis* infections (exclusively from Africa or linked to refugee status), with a caveat that both ancient and modern diversity may be significantly undersampled. A divergence date from *B. duttonii* Ly dating to at least the Neolithic implicated in our temporal analysis is supported by a preprint study providing a low-coverage observation of a 4,600-year-old Late Neolithic individual from Denmark likely infected by *B. recurrentis*^29^ (**Figure 2B**).

To validate the Bayesian inference of a relatively recent divergence of *B. recurrentis* from the shared ancestor with *B. duttonii* Ly, we also performed an additional analysis. Here, we identified SNPs where *B. crocidurae* DOU and *B. duttonii* Ly both had an alternative variant to all modern *B. recurrentis* genomes, which we can interpret as new mutations occurring on the lineage leading to the modern *B. recurrentis* clade since the divergence **(Methods)**. This approach also has the advantage of excluding any impact of sequence errors specific to the ancient genomes. If divergence occurred approximately 5,000 years ago, we could expect the ancient individuals to have accumulated these mutations in an approximately clock-like manner, with e.g. the ∼700-year old medieval genome, C10976 Poulton, having at most 1-(700/5000)=86% of these derived mutations, and an Iron Age ∼2,200-year-old genome having at most 57%. Indeed, these expectations match what we observe in the empirical data (**Table S4**), and we find an intercept in linear regression of ∼6,100 years ago, overlapping with our estimates following Bayesian tip-dating calibration when C13361 Fishmonger’s is included. We note that this number is expected to slightly overestimate the true divergence because the TMRCA of *B. recurrentis* A1 and the ancient genome will always be slightly older than the age of the sample.

### Patterns of pan-genome diversity and genome reduction in *B. recurrentis*

We used a pan-genome approach to establish patterns of gene content across *Borrelia* (**Methods**) and noted variability in the number of genes per species congruent with previous observations. We identified that *B. recurrentis* contains ∼25% fewer genes than *B. duttonii,* with the least variance in gene content in the pan-genome of any relapsing fever species in the genus, consistent with its hypothesised niche constraint^30^ (**Figure S8, Figure S9**). This result is replicated when directly aligning to the *B. duttonii* Ly reference genome. We estimate that during the medieval period, the high coverage C10976 Poulton genome covered ∼82% of the *B. duttonii* reference (the previously published genome OSL9 covered ∼75%, which may be potentially underestimated due to lower coverage) and the older Iron Age genome, C10416 Wetwang Slack, covered ∼81%. This demonstrates that almost all of the reductionist evolution seen in *B. recurrentis* had already occurred ∼2,000 years ago.

We assessed the contribution of plasmid carriage relative to *B. duttonii*, *B. crocidurae* and contemporary *B. recurrentis* (**Figure 3**). We identify three plasmids (pl26, pl27, pl28), described in *B. duttonii* Ly, that are present in the Iron Age genomes—authenticated using coverage, cytosine deamination patterns and distribution of mismatches—but are absent or at substantially lower coverage in medieval and present-day genomes (**Figure S10, Table S5**). We therefore suggest at least partial plasmid loss events, or loss of significant plasmid-borne elements, ∼2,000-700 years ago, between the Iron Age and medieval lineages. Overall, by the medieval period, *B. recurrentis* harboured the full suite of plasmids observed in currently sampled infections. In contrast, the Iron Age genome shows only partial coverage of *B. recurrentis* plasmid pl53, which is present in medieval and modern genomes, suggesting that the complete plasmid gene complement was acquired after ∼2,000 years ago (**Table 1**, **Figure 3**).

**Figure 3.**
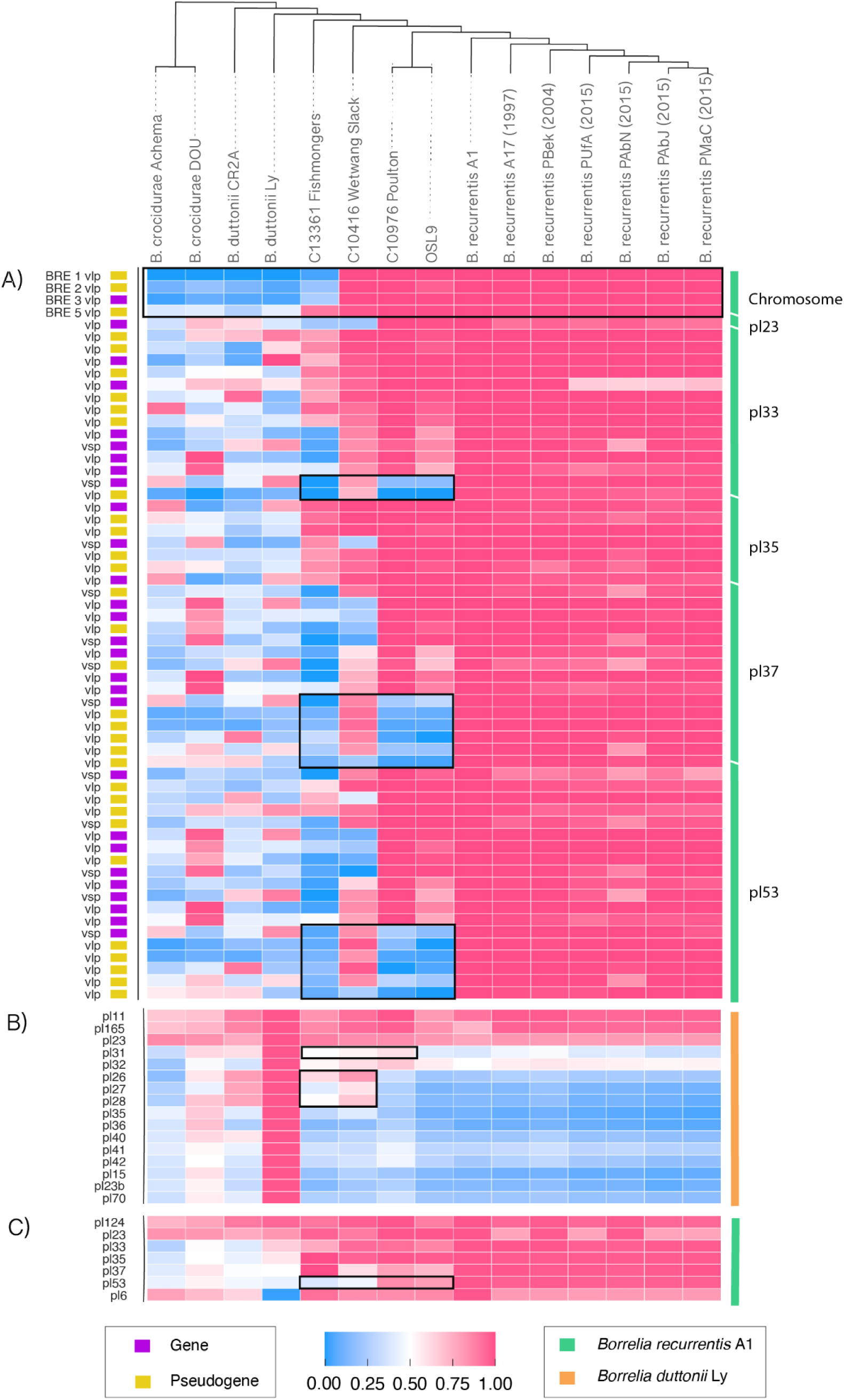
Gene losses/gains across variable major proteins (vmp), *B. duttonii* Ly and *B. recurrentis A1* plasmids. Ancient and modern genomes were aligned to single reference *B. recurrentis* (green) or *B. duttonii* (orange) (**Methods**). Regions of interest highlighted in the text are outlined with a black box. Cladogram provides the relationship between different genomes based on a SNP phylogeny. **A)** Normalised coverage across the variable major proteins on the *B. recurrentis* A1 chromosome and plasmids (pl), using *BEDTools* v2.29.2. Coordinates of the vmp genes and whether they are classified as genes (yellow) or pseudogenes (purple) were provided in Guellil et al^17^ using previously annotated genomes from the NCBI database. **B)** Breadth of coverage for *B. duttonii* Ly plasmids using SAMTools v1.3.1 with a mapping quality of Q1. **C)** Breadth of coverage for *B. recurrentis* A1 plasmids using *SAMTools v1.3.1* with a mapping quality of Q1.

We next assessed gene variation, agnostic to chromosomal or plasmid affiliation, captured by our ancient *B. recurrentis* genomes, given that they may represent possible intermediates on the trajectory towards specialisation. First, we assessed the full gene repertoire across all *B. recurrentis, B. duttonii and B. crocidurae* species. A total of 3035 unique genes were identified, defining the pan-genome, which were then subjected to presence/absence assessment, filtering and annotation (**Methods**). Given hypotheses surrounding genome reduction and plasmid stability, we particularly noted two genes in the pan-genome implicated in plasmid segregation and partitioning, *Soj* and *ParA* (annotated as *Soj_1* and *ParA_1*), which showed temporal patterning across our dataset (**Table S6**, **Figure S11**). *Soj_1* is present in both *B. crocidurae* and *B. duttonii*, and the ancient Iron Age *B. recurrentis* genomes of sufficient coverage for consideration, but absent in medieval and present-day data. Assuming parsimony, this suggests gene loss on the branch leading to the descendant medieval and contemporary clades (we estimate phylogenetically, based on current data, between 2,326-1,115 years ago; 95% HPD). In contrast, *ParA*, an ortholog of the *Soj* gene^31^ and also implicated in plasmid segregation, is present solely in the medieval and modern-day genomes. A parsimonious explanation is the acquisition of this gene between the Iron Age and the medieval period. Exploration of the genomic neighbourhoods of both the *ParA_1* and *Soj_1* genes (**Methods**) further supports that these genes are today localised on different backgrounds, with *Soj_1* found on the *B. duttonii* Ly chromosome and *ParA_1* on the pl53 *B. recurrentis* plasmid.

### Temporal variation in functional genes

*Borrelia* relapsing fevers use a suite of predominantly plasmid-encoded antigenic phase variation as a mechanism of immune evasion. This is mediated by variable large proteins (vlp) and variable short proteins (vsp), together known as the variable major proteins (vmp). Antigenic variation of these surface-exposed lipoproteins likely plays an important role in evading host-acquired immunity, allowing for the bacteria to persist within its host population. However, it is unclear how stable this mechanism has been through evolutionary time^32–34^. Of the chromosomal vmp genes, we found that the medieval lineage, comprising C10976 Poulton and OSL9, had similar vmp profiles, both to each other and to present-day *B. recurrentis,* with three of the four chromosomal-borne vmps characteristic of present-day *B. recurrenti*s having been gained by medieval times (**Figure 3**). However, two of these vlp genes have been identified as pseudogenes in modern *B. recurrentis* genomes, and so, despite being gained in *B. recurrentis*, these are believed to be functionally redundant in modern genomes.

When evaluating vmp profiles over the *B. recurrentis* plasmids, we noted the absence of a number of vmp genes in the medieval genome, many of which are pseudogenes. This was seen in particular at the 3’ ends of the pl33, pl37 and pl53 plasmids (**Figure 3**); an observation also seen in the previously published OSL9 genome from the same time period^17^. Interestingly, these regions (5’ pl35, pl37 and pl53) are mostly present in the Iron Age sample C10416 Wetwang Slack but are absent in our basal Iron Age sample C13361 Fishmonger’s. Due to the lower overall coverage of the Fishmonger’s genome, the exact gene complement of plasmid-borne vmp genes is difficult to formally assess. However, this observation is consistent with the potential for interspecies variability in vlp profiles, as is seen in its tick-borne relatives (*B. crocidurae* and *B. duttonii*).

Evaluating other hallmarks of infective behaviour^3^, we note that BDU 1, a p35-like antigen implicated in fibronectin binding in *Borrelia burgdorferi* (Lyme disease)^3,35^, is absent in *B. recurrentis*. This outer membrane protein is also absent across our Iron Age and medieval observations supporting an early loss following the split with *B. duttonii Ly (***Figure S11)**. A similar pattern was seen for several further outer membrane proteins, including those involved in host complement system inactivation (BDU 2, BDU 3, BDU 5)^33^, which are absent in *B. recurrentis* from ∼2,000 years ago by our earliest observation. We do however observe some temporal variation which suggests an ongoing process of genome adaptation, for example, the aforementioned loss of *Soj* (annotated as BDU 429) and truncation of the uncharacterised protein, BDU 430, in the modern *B. recurrentis* genome. These genes are also absent/truncated in the medieval genomes, but they are present in the Iron Age genomes.

Finally, we assessed SNPs and indels of functional relevance (**Figure S12**)^3^. For example, the *oppA-1* gene is a pseudogene in *B. recurrentis* A1 due to an in-frame stop mutation; but in *B. burgdorferi*, it is shown to play an essential role in metabolic function as well as survival in different environments^36^. As with the previously reported medieval genome from OSL9^17^, this gene is found in its ancestral form in all of our ancient samples. This suggests the inactivation of *oppA-1* occurred relatively recently, we estimate within the last ∼1,115 years (**Figure 2**). Similarly, the *smf* gene and the *mutS* gene show an in-frame stop mutation and a frameshift mutation in present-day *B. recurrentis*. All ancient genomes with data at this locus show the in-frame stop mutation in the *smf* gene, with the higher coverage genomes (C10416 Wetwang Slack and C10976 Poulton) also supporting the presence of the *mutS* frameshift mutation. Conversely, the *recA* gene is still functional in the high-coverage Iron Age genomes. Hence, the true recombination efficiency is unknown for these genomes even if we find little detectable signal of recombination in our genomics analysis (**Figure S5**).

## Discussion

Here we reconstruct the complex evolutionary history of *B. recurrentis* by retrieving and analysing four ancient *Borrelia* genomes from Britain across a ∼1,500-year time span. Our work confirms the presence of the pathogen in Europe during both the Iron Age and the later medieval periods, extending the high-coverage *B. recurrentis* whole-genomes by approximately 1,600 years. This detection adds markedly to existing data from the species, with only eight genomes (seven contemporary and one medieval) available prior to this study. While it is unclear whether there is any link between our detected infections and historically-attested outbreaks in Britain, we note that the high coverage recovery of genomes achieved here suggests that the individuals studied likely died from acute infections with high levels of bacteremia. The likely high false negative rates of infectious disease detection in ancient remains means that it is not possible to link these results with absolute rates of disease amongst past populations more generally. Without antibiotic treatment LBRF infections are fatal in 10-40% of cases^37^, however it is uncertain to what extent this figure would be applicable to ancient cases of the same disease which differed genetically from the modern forms and was also operating in variable cultural and environmental contexts.

In the context of published datasets, our work confirms the existence of a closely related medieval phylogenetic clade that existed from at least 600 years ago and spanned Britain and the Scandinavian peninsula. In addition, we recover previously unknown ∼2,000-year-old basal lineages from Iron Age Britain. The phylogenetic placements of the Iron Age genomes may suggest that multiple lineages of *B. recurrentis* existed at this time. Harnessing this temporal structure, we find support for a relatively recent divergence of sampled *B. recurrentis* strains from the closest relative *B. duttonii* Ly within the last ∼8,000 years (across models and datasets). While we caution that a more complete time series could improve these calibrations and divergence estimates, the evidence is consistent with a Late Neolithic/Early Bronze Age emergence of the agent of LBRF. This could have been linked to factors such as changes in human lifestyles which were more favourable to the human body louse vector. Good examples of this would be the gradually increasing levels of sedentism and contact with domesticates during the development of agriculture, as well as the emergence of densely-occupied mega-settlements in regions of Eastern Europe specifically^38^.

To date, no *B. recurrentis* whole genomes have been identified earlier than these estimates. Sikora et al^29^ report 31 observations of *B. recurrentis* in the ancient skeletal record in an unpublished preprint. These findings are based on the assignment of sequencing reads from human skeletons, mostly from Eurasia (**Figure 2B and Figure S13**), all of which are lower than 0.17-fold genome coverage. The earliest recovered observation is from Denmark dating to

∼4,600BP (RISE61; 0.003X)^29,39^. Due to the low coverage of these *B. recurrentis* observations through time, it was not possible to incorporate them into our phylogenetics analysis. However, further sequencing of these or other ancient samples may prove vital in establishing the spatio-temporal timeline of infections, as well as to potentially uncover previously unseen genomic diversity. Contemporary infections are also undersampled, with limited available genomic data mostly linked to cases in East Africa or associated with migrants on their journey to Europe^37^. As such, it is challenging to disentangle many of our observations of ancient strains from the expectation of population structure between African and European strains. Further data from different time periods and geographic regions of the world may result in our temporal estimates being amended. This would be required to assess any connection between the population structure of the bacteria and the mobility of its human hosts.

Our estimates over the non-recombining proportion of the genome indicate a recent timeline over which presently sampled *B. recurrentis* diverged from its closest sequenced relative, *B. duttonii* Ly. We note that some have suggested that *B. recurrentis* is a degraded form of *B. duttonii* and hence the lines between species demarcations may well have been blurred in deeper history. Aside from undersampling, we must also consider the possibility of rate variation in the history of *B. recurrentis*. While a relaxed clock model was less well supported by our Bayesian phylogenetic analyses, it is plausible that ecological influences on mutation rate, as observed in other bacterial pathogens, may have played a complex role which is difficult to capture using our temporal reconstruction approach^40,41^.

Nonetheless, our work indicates the need for rapid genome decay in order for *B. recurrentis* to exhibit ∼20% reduction in its genome relative to *B. duttonii* Ly within ∼8,000 years; an evolutionary behaviour which has been linked to specialism towards the human body louse vector and potentially resulting in enhanced pathogenicity^42^. We observe that a striking amount of this decay had already occurred by the time of our oldest sampled infection ∼2,000 years ago. It has been suggested that this accelerated evolution, particularly in light of the slow global mutation rate, may have been supported by plasticity in the wider accessory genome of *B. recurrentis*. It remains unclear precisely to what extent the genome size differs between the two species due to the reductive evolution in *B. recurrentis*, as we also observe some unique gains in the *B. duttonii* Ly genome. However, aided by the high coverage Iron Age and medieval *B. recurrentis* genomes, we can demonstrate that at least some of the decay towards the extant *B. recurrentis* genome was ongoing between these periods. In particular, we identify the partial loss of three plasmids (or extensive plasmid-borne elements) between *B. duttonii* and our Iron Age samples and later medieval and contemporary *B. recurrentis* strains; we estimate that this event most likely occurred between 2,326 and 1,115 years ago.

The extent to which such loss events may still be ongoing is unclear, though it has been suggested that disruption in plasmid partitioning genes relative to *B. duttonii* homologs may indicate a degree of an ongoing reductive process^3^. Indeed, we note an interesting temporal pattern of major chromosomal and plasmid partitioning genes *Soj* and *ParA*, best described for their role in *Bacillus subtilis* and *Escherichia coli*^43^. While it has previously been reported that *B. recurrentis* lacks a chromosomal *Soj* homologue^3^, using a pan-genomics approach we identify that *Soj* was retained until at least the Iron Age. The homologue was then lost by the earliest of our two medieval observations, where we simultaneously reconstruct the gain of *ParA*, a distant homolog of *Soj*. Such observations highlight the fluidity of the process of genome reduction, with the suggestion of necessary acquisition of some functionally relevant genes as a likely outcome of large-scale plasmid loss events. Another plausible contributor to the pattern of genome decay is the loss of DNA repair mechanisms, with the inactivation of genes such as *recA* resulting in the bacteria becoming dependent on its human/vector hosts^8^. Our data suggest that the loss of *recA* as a DNA repair mechanism may also be a reasonably recent event, given that we find the *recA* gene is still functional in our Iron Age observation. It is significant to note that both *smf* and *mutS*, also implicated in DNA repair, are disrupted across our ancient samples, supporting the importance of this mechanism in the wider propensity for genome loss.

The transition to a host specialised pathogen from a more generalist ancestor will also have exerted a selective pressure on the bacteria. Within *Borrelia*, the vmp and vmp-like genes offer an important mechanism to allow persistence and resurgence of relapsing fevers, with antigenic variation during infection through a process of silent vmp genes being transferred to the expression locus, leading to the generation of new surface protein variants^32–34^. Our work supports some temporal variability within *B. recurrentis*, particularly in the vmp genes located at the 3’ end of the pl33, pl37 and pl53 plasmids, which we observe as absent in medieval samples. While this may represent a change in antigenic behaviour, it is also plausible that the pl33, pl37 and pl53 plasmids are shorter or subject to genomic rearrangements in these strains. This observation was also suggested by Guellil and colleagues who detected a similar patterning in the only other ancient *B. recurrentis* full genome published prior to this study^17^. We also note far richer diversity in vmp profiles in non-*recurrentis* species, indicating that the extent of antigenic plasticity may have been very different prior to host specialisation.

Together we highlight how ancient microbial DNA can be used to enhance our understanding of the age and diversity of significant but understudied pathogens. Our work highlights the value of temporal data in pinpointing the timing and patterning of the process of host/vector specialisation, supporting a prevailing background of accelerated genome reduction, notwithstanding more recent key instances of gene gains and losses. We can, however, only speculate on whether these ancient bacteria from Britain were adapted to be a louse-borne or tick-borne form of relapsing fever. Additional work is required to build a mechanistic understanding of the genomic basis for each vector niche.

## Acknowledgements

We thank the Advanced Sequencing Facility at the Francis Crick Institute for technical support. This work was supported by the Vallee Foundation, the Wellcome Trust (217223/Z/19/Z), and Francis Crick Institute core funding (FC001595) from Cancer Research UK, the UK Medical Research Council, and the Wellcome Trust. P. Skoglund was supported by the European Molecular Biology Organisation, and the European Research Council (grant no. 852558). L.vD was supported by a UKRI Future Leaders Fellowship MR/X034828/1 and European Union END-VoC consortium grant (agreement no. 101046314). P. Swali was supported by the Francis Crick Institute core funding (FC001595 to P. S.) and a UKRI Future Leaders Fellowship grant (MR/X034828/1 to L.vD.). This research received funding from the European Research Council (ERC) under the European Union’s Horizon 2020 research and innovation programme (grant agreement No. 834087; the COMMIOS Project to I.A.) L.S. was supported by a Sir Henry Wellcome Fellowship (220457/Z/20/Z). We are grateful to the University of Bristol Spelaeological Society, the University of Bradford, Hull and East Riding Museum and Canterbury Archaeological Trust, Liverpool John Moores University, and the Poulton Archaeological Trust for allowing access to collections and facilitating sampling. We would also like to thank Meriam Guellil and Marina Escalera for help and advice in the analysis.

## Author contributions

Conceptualisation: P. Swali., L.vD., P. Skoglund.; Software: P. Skoglund.; Data processing and curation: P. Swali., C.B., A.G.; Formal analysis: P. Swali., L.vD.; Visualisation: P. Swali.; Investigation: P. Swali., T.B., C.T., J.M., K.A., C.B., A.G., I.G., M. Kelly., J.P., M.S., L.S., F.T., M.W., L.vD., P. Skoglund.; Resources: M.B., A.B., J.B., L.B., R.C., G.M., R.M., J.I., M.King., F.P., J.P., A.T., S.V., L.W., K.C., I.A.; Supervision: L.vD., P. Skoglund.; Writing—original draft: P. Swali., L.vD., P. Skoglund.; Writing – review & editing: P. Swali., T.B., C.T., L.vD., P. Skoglund.

## Data Availability

All sequence data will be available in the European Nucleotide Archive upon publication.

## Code availability

Code to reconstruct a consensus sequence using strand-aware removal of cytosine-deamination-derived errors is available at (https://github.com/pontussk/mpileup2consensus.py/).

## Methods

### Archaeological Context

Wetwang Slack is an Iron Age Arras-style cemetery in East Yorkshire dating to 2300-2100 years ago (300-100 BCE). Sample C10416 was taken from a left mandibular third molar from Burial 240. Fishmonger’s Swallet is a cave site in South Gloucestershire. Sample C13361 was taken from a left first molar in a disarticulated human mandible (G10-1.4) that has been radiocarbon dated to 2185-2033 years ago (162 cal. BCE - 10 cal. CE; 2063±28 BP, BRAMS-5059^20^). Sample C11907 was taken from a maxillary left third molar from an ancient cranium (CW29) held by the Canterbury Archaeological Trust (CAT). The provenance of CW29 is uncertain but based on the collections held at CAT the cranium must be from a site in Canterbury or the surrounding region and dates to 2,000-700 years ago (0-1,300 CE). Sample C10976 was taken from a maxillary canine of an adult male (Sk 435) buried in a cemetery associated with a medieval chapel at Poulton, Cheshire. Radiocarbon dates from Sk 435 and other parts of the cemetery indicate that Sk 435 dates to 733-633 years ago (1290-1390 cal. CE, 646±14 BP, Wk 52986^21^). For more information on the archaeological context see **Supplementary Note S1**.

### Sampling, DNA extraction and library preparation

One tooth from each individual included in this study was sampled and processed in a dedicated cleanroom facility at the Francis Crick Institute. An EV410-230 EMAX Evolution Dentistry drill was used to clean the surface of the tooth and both the cementum and multiple fractions of the dentine were sampled, resulting in ∼11-35 mg of powder from the dentine. 300 µl (<10 mg of powder), 600 µl (10-25 mg) or 1000 µl (>25 mg of powder) of extraction buffer (0.5 EDTA pH 8.0, 0.05% Tween-20, 0.25 mg/ml Proteinase K^44^) was added to the dentine powders and incubated for 24 hours at 37℃. They were then centrifuged for 2 minutes at 13,200 rpm in a table centrifuge and 140 µl of the supernatant was transferred into LVL tubes for automated extraction on an Agilent Bravo Workstation^45^. Extracts were turned into single-stranded DNA libraries^46^, then double-indexed^47^ and underwent paired-end sequencing with a 2×100 paired-end read configuration on the Illumina HiSeq4000 or NextSeq500 and Novaseq platform (**Table 1** for sequencing effort per library). All samples were processed alongside negative extraction controls as well as positive and negative library controls.

One library, C10416 from Wetwang Slack, underwent size selection to remove fragments shorter than 35 bp and longer than 150 bp, as in Gansauge et al. 2020^46^. Specifically, 100 ng of the initial library was biotinylated and streptavidin beads were used to isolate the non-biotinylated strand and obtain a single-stranded library. This sample was then loaded on a denaturing polyacrylamide gel along with 35 bp and 150 bp insert markers, and fragments within the desired sequence length were physically excised and eluted from the gel, after overnight incubation. The resulting size-selected libraries were further amplified and sequenced on the Illumina NovaSeq.

### Bioinformatic Processing, Metagenomic Screening and Authentication

Samples were initially processed via the nf-core/eager v2 pipeline^48^. First, adapters were removed, paired-end reads were merged and bases with a quality below 20 were trimmed using AdapterRemoval v2^49^ with –trimns –trimqualities –collapse –minadapteroverlap 1 and –preserve5p. Merged reads with a minimum length of 35 bp were mapped to the hs37d5 human reference genome with Burrows-Wheeler Aligner (BWA-0.7.17 aln)^50^ using the following parameters “*-l 16500 -n 0.01*”^18,51^. We then analysed sequences that did not align successfully to the human genome using Kraken2^52^ and identified individuals as putatively positive for *B. recurrentis* by assessing an excess number of observed *k*-mers (sequence matches).

These libraries were subsequently aligned to the *B. recurrentis* A1 reference genome (chromosome and plasmids; GCF_000019705.1) using BWA-0.7.17 aln^50^ parameters "*-l 16500 -n 0.01 -o 2*". Duplicates were removed by keeping only the first sequence out of any set of sequences with the same start position and length (https://github.com/pontussk/samremovedup). We assessed the authenticity of the final set of sequences using the following criteria^24^: i) the observation of postmortem damage, ii) the number of sequences being negatively correlated with edit distance from the reference genome, and iii) an unimodal fragment length distribution via DamageProfiler^53^ iv) even breadth of coverage across the *B. recurrentis* A1 reference genome using SAMTools v1.3.1 *depth*^54^. Additionally, these libraries were aligned to the *B. duttonii Ly* and the *B. crocidurae DOU* reference genome (NC_011229.1 and NZ_CP004267.1 respectively) and their edit distance distributions were compared (**Figure S2**). Screening libraries that passed these authentication criteria, were taken forward for further shotgun sequencing. For the final BAM files, we merged shotgun BAM files using SAMTools *merge* resulting in a final chromosome coverage of 29.4X, 11.2X, 3.5X and 0.8X coverage for C10976 Poulton, C10416 Wetwang Slack, C13361 Fishmonger’s and C11907 Canterbury, respectively when aligned to *B. recurrentis* A1 (**Table 1, Figure S1**).

### Dataset Curation

All published genomes used in this study are listed in **Table S1**. This includes all modern genome assemblies available from NCBI (accessed April 2024). In addition, we *de-novo* assembled six modern genomes available on the SRA linked to BioProject PRJNA378726 using UniCycler v0.50 in short-read only mode, recovering high quality assemblies (N50>700,00 in all cases). The previously sequenced ancient *B. recurrentis* genome OSL9^17^ was downloaded from ENA following which adapters were removed and reads were merged using AdapterRemoval2 and processed identically to the described method for individuals in this study (**Methods, Bioinformatic Processing**).

### Alignment Approaches and Phylogenetic Reconstruction

Two approaches were used to construct an alignment for phylogenetic inference. The first used a reference mapped approach to the *B. recurrentis* A1 reference genome (as described above). The second was to construct a core gene reference alignment, built using an assessment of gene content in modern strains. For the latter, we initially applied Panaroo v1.1.2 on the 11 (seven *B. recurrentis,* two *B. duttonii* and two *B. crocidurae)* modern RF genomes and assemblies available specifying the *–core* flag in relaxed *-mode* to obtain a core alignment of genes featuring in 99% of considered genes. We then aligned all genomes (ancient and modern) to the core gene sequence using BWA-0.7.17 *aln* parameters "*-l 16500 -n 0.01 -o 2*", and processed to remove duplicates by keeping only the first sequence in case multiple sequences had the same start and end positions (https://github.com/pontussk/samremovedup). For both approaches, for published assemblies, where short-reads were not available, we applied SeqKit to generate pseudo-reads for which the mapping pipeline could be applied.

In both cases, modern genomes were converted to fasta sequences keeping all bases with a coverage of 1 using HTSBOX^55^. For the ancient genomes, we computed Base Alignment Qualities using SAMTools "*mpileup -E*", restricted to a minimum phred-scaled mapping quality of 30 and base quality 30 using SAMTools v1.3.1. The MD field was modified to record mismatches to the reference using SAMtools *calmd*. Relative to the A1 reference genome and core genome alignment, C to T transition mutations on the forward strand and G to A transition mutations on the reverse strand were masked to correct for the possible effects of cytosine deamination in the single-stranded ancient DNA sequences (https://github.com/pontussk/mpileup2consensus.py/blob/main/mpileup2consensusfasta.py). Additionally, using this tool we filtered out all heterozygous base calls and only kept sites with a minimum coverage of 3 calls per site, resulting in a filtered fasta for each ancient genome.

Ancient and modern Fastas were then concatenated and polymorphic positions were identified for the initial maximum likelihood phylogeny. Using https://github.com/pontussk/fasta_nomissing.py, polymorphic positions with a threshold of maximum missingness of 20% per site and a maximum missingness of 20% per genome were implemented resulting in 4,200 sites when aligned to the core-genome (**Figure S3**) and taken forward for maximum likelihood phylogenetic reconstruction in IQ-TREE v.1.6.12. Due to these thresholds, C11907 Canterbury was excluded from most downstream analysis. We implemented model testing using the ModelFinder in IQ-TREE^56^, which suggested a TIM+F+ASC as the best-fit model according to the Bayesian information Criterion. We implemented 1000 rapid bootstrap replicates and rooted the maximum likelihood phylogeny in FigTree using *B. duttonii Ly* as an outgroup.

### Recombination and Temporal analysis

Using the initial SNP tree and the whole core-genome alignment, ClonalFrameML v1.13 was applied to detect homoplasies and putative recombination tracts. Identified recombinant tracts were masked from the alignment **(Figure S5)**.

The recombination pruned core genome alignments and corresponding phylogenies were inspected for signatures of temporal evaluation using the *roototip()* function implemented in BactDating^27^, evaluating empirical significance following 10,000 randomisations of the sampling dates. Temporality was assessed for a dataset including and excluding the C13361 Fishmonger’s sample due to its low coverage, and using phylogenies built on alignments with and without the inclusion of transitions. In all cases we obtained a highly significant temporal regression (**Table S2**).

Resulting alignments were subsequently taken forward for formal Bayesian tip-dating calibration implemented in the BEAST2 workflow^28^. In all cases, variant positions were considered (with corresponding correction for the base composition of invariant sites) with the prior on the tip dates corresponding to the date of sample collection, or, where a range of dates was given (either in contemporary samples or corresponding to radiocarbon dates) the mean estimate was used as an initial prior. Following evaluation of possible substitution models in BModelTest, in all cases a GTR substitution model was best supported and selected for further analysis. BEAST2 was run assuming either of a strict or relaxed (exponential distribution) molecular clock exploring three distinct priors for the demographic model: coalescent constant, coalescent exponential and coalescent bayesian skyline, specifying 50 million chains sampling every 1000. Resulting chains were inspected for convergence in TRACER, requiring an effective sampling space (ESS) of >200, with resulting posterior estimates extracted following discarding the first 10% of chains as burn-in. All runs exhibited a significant difference between the posteriors obtained when sampling from the prior (data absent model). Finally, model fit was assessed using the path-sampling model, requiring 100 steps over 250,000 chains, to establish marginal likelihoods for the models and corresponding Bayes Factors. All results are provided in **Table S3** with posterior distributions available in **Figures S7** and **Figures S8**.

We also took an alternative approach to approximating the divergence time, which excludes external branches private to ancient genomes, and thus removes any potential impact of sequence errors unique to the ancient genomes. To identify ancestral and derived variants separating modern *B. recurrentis* from the common ancestor with *B. duttonii* Ly (**Table S4**), we used the core-genome aligned multifasta containing the ancient genomes which had been filtered using mpileup2consensusfasta.py and all modern *B. recurrentis*, *B. duttonii* Ly and *B. crocidurae* DOU genomes and generated a VCF using SNP-sites^57^. We then filtered the VCF to keep only SNPs where all modern *B. recurrentis* genomes matched the *B. recurrentis* A1 genome, and ancestral where both *B. duttonii* and *B. crocidurae* were identical to each other but differed from the *B. recurrentis* variant. We calculated the number of variants that were missing, ancestral or derived in our ancient genomes (**Table S4**) and used the % derived and the approximate date of the ancient genomes to identify an approximate TMRCA based on the intercept of linear regression.

### Pangenome Analysis and Evaluation of Gene Content

We constructed a full pan-genome given diversity in modern *B. recurrentis*, *B. duttonii* and *B. crocidurae* genomes (**Table S1**). To do so we initially applied Prokka v1.12^58^ to generate gene annotations for each modern genome. Panaroo v1.1.2 with the -*pan* flag in relaxed mode and otherwise default thresholds was implemented and then used to identify gene clusters and create a list of genes present in all of the given modern genomes. We identified 14,475 genes and gene clusters across all modern relapsing fever *Borrelia* species, which were subsequently filtered for pseudogenes or genes of unusual lengths. Among the modern *B. recurrentis* genomes, Panaroo identified ∼1,100 of these genes as present.

To identify the presence and absence of genes in the pan-genome we aligned simulated short-read modern FASTQs using BWA-0.7.17 *mem* with the parameters “*-B 40 -O 60 -E 10 -L100*” to the pangenome and the ancient FASTQs using BWA *aln* with the parameters "*-l 16500 -n 0.01 -o 2*". We then used the pangenome gene clusters. This allowed us to set an arbitrary threshold to identify genes as present required a minimum threshold of 70% coverage and below 30% coverage for absence. In addition we only kept genes which showed phylogenetically congruent patterning, resulting in consideration of 71 well resolved genes of interest **(Table S6**). We performed *in silico* functional predictions for these genes (**Methods**) using the EggNOGG-Mapper web server (http://eggnog-mapper.embl.de/) and InterproScan^59^, both with default parameters. Putative gene functions were inferred from the annotations provided by both tools **(Table S7).** The gene neighbourhood of identified hits, including *Soj_1* and *ParA*, were assessed using the Panaroo pan-genome network graphs, visualised in Cytoscape v3.10.2. In this case, gene homology was obtained by translating the nucleotide sequence of *Soj_1* and *ParA* to amino acids before aligning in MUSCLE and calculating pairwise identity.

### Visual inspection of SNPs and Indels

An additional reference based analysis was also carried out, aligning all genomes to *B. recurrentis* A1 and *B. duttonii* Ly. Here, all ancient and simulated short-read modern genomes were aligned to each of the aforementioned references using the described BWA aln pipeline for ancient samples and BWA *mem* for the modern genomes (specifying the same parameters as when aligning to the pangenome). After the removal of duplicates, we assessed the coverage across both chromosomes, plasmids, and previously reported genes, and manually inspected previously reported SNP mutations using Intergrated Genome Viewer (IGV) (**Figure S12**). Additionally, we assessed regions of missingness by identifying regions with a gap in coverage larger than 500bp in the ancient genomes using BEDTools^60^ when aligned to each of the *B. recurrentis* A1 reference genome and the *B. duttonii* Ly reference genome (**Figure S15**, **Table S8**). We noted some marked drops in coverage consistent with genomic deletions or rearrangements. As an example, despite the low coverage of C11907 Canterbury, we could confirm a unique 26.1kb deletion when aligned to both colinear plasmid pl124 in *B. recurrentis* A1 and pl165 in *B. duttonii* Ly (**Figure 1B, Figure S14, Table S8**). The pl124 plasmid in *B. recurrentis* A1 has been previously observed to exhibit a ∼40kb deletion in comparison to the *B. duttonii* plasmid^3^. Marosevic et al^8^ identified this 40 kb region to also be present in eight modern-day *B. recurrentis* genomes from East Africa, suggesting this deletion may be a recent event, or potentially the product of an incomplete assembly of the A1 reference genome plasmid. We find this 40kb region is also present in all of our ancient *B. recurrentis* observations, consistent with these proposed scenarios.

## Supplemental Information

### Supplementary Note S1

#### Archaeological Context

##### Wetwang Slack (Ian Armit)

The individual with LBRF is Burial 240 at Wetwang Slack, a square-ditched barrow cemetery in East Yorkshire associated with the ‘Arras Culture’ dating to 300-100 BCE (Armit 2021, Dent 2010). The human remains are curated at the University of Bradford on loan from the Hull and East Riding Museum. This individual, identified osteologically as an adult female, was buried without any accompanying grave goods under a square barrow. The lack of grave goods is not especially unusual at Wetwang Slack, even for primary barrow burials such as this. This individual has been identified as the mother or daughter (order currently unknown) of another adult female (Burial 303), and a second-degree relative of a third adult female (Burial 270), both also primary burials under square barrows. This suggests that she was likely a locally-born member of the community and, given that she was a primary rather than secondary burial, she was probably of reasonably high status (although not in any major hierarchical sense given the numbers of barrows present at Wetwang Slack).

##### Fishmonger’s (Adelle Bricking, Graham Mullan, Linda Wilson)

Fishmonger’s Swallet is a small stream sink located near the village of Alveston in the Vale of Berkeley, South Gloucestershire. Initial clearing of the swallet was undertaken by Clive Grace, a local fishmonger, from whom the site derives its name. Subsequently, David Hardwick and the Hades Caving Club assumed control of the dig, revealing a substantial quantity of disarticulated human and animal bones, particularly canids, concentrated within an area designated the Bone Idle Chamber (Hardwick 2022). This discovery attracted the attention of the Channel 4 programme Time Team and a three-day archaeological investigation conducted by the team uncovered additional human and animal bone fragments, along with evidence of post holes on the surface near the swallet (Horton 2022).

Post-excavation analysis of the human remains by Cox and Loe (2022) showed a minimum number of five individuals, but possibly six adult females and five adult males represented by the assemblage. The degree of fragmentation and environmental staining is limiting for macroscopic observations, nevertheless some evidence of pathology (Paget’s disease, degenerative joint disease and abscesses) and perimortem trauma was seen on some elements (Cox and Loe 2022).

It is important to note that a small creek flows into the swallet from the surface, carrying modern debris down a short vertical shaft into the Bone Idle Chamber. The co-mingling of modern and archaeological material means it is impossible to interpret the depositional history of the site stratigraphically, so radiocarbon dating is essential. A total of sixteen radiocarbon dates have been obtained from skeletal material and recent programme comprising seven human (BRAMS-5057, BRAMS-5058, BRAMS-5059, BRAMS-5060) and six canid (BRAMS-5061, BRAMS-5062, BRAMS-5063, BRAMS-6671, BRAMS-6672, BRAMS-6673) bone have produced dates centred on the later Iron Age (Bricking et al. 2022, Bricking and Mullan forthcoming).

##### Canterbury (Adelina Teoaca)

Sample C11907 comprised dentine powder taken from a left maxillary third molar from a complete cranium (CW 29) held in the archives of Canterbury Archaeological Trust (CAT) in Canterbury, Kent. The provenance and date of cranium are uncertain but the scope of CATs archive both today and historically mean that it must come from eastern Kent and date to 2000-700 years ago (0-1300 cal. CE).

##### Poulton (Kevin Cootes)

Sk 435 was recovered during the excavation of a medieval burial ground and associated chapel in rural Poulton, Cheshire, England. The site was used for interment of individuals from a small farming community, with historical documents and radiocarbon dates indicating use between the 13th and 16th centuries CE^21,61^. The remains are curated at Liverpool John Moores University as part of their teaching and research collection. Sk 435 was identified osteologically as a mature adult of undetermined sex, who was approximately 40-44 years old at the time of death. The body was laid out in typical medieval fashion, extended in a supine position with west–east orientation. Neither grave goods nor a burial marker were present. Continual re-use of the cemetery resulted in the individual’s legs and portions of the pelvis being removed by later interments. Radiocarbon dating of the remains provided a date of 1290-1390 cal. CE at 2 δ (Wk52986. 646 ± 14BP).

## Supplementary Figures

**Figure S1.**
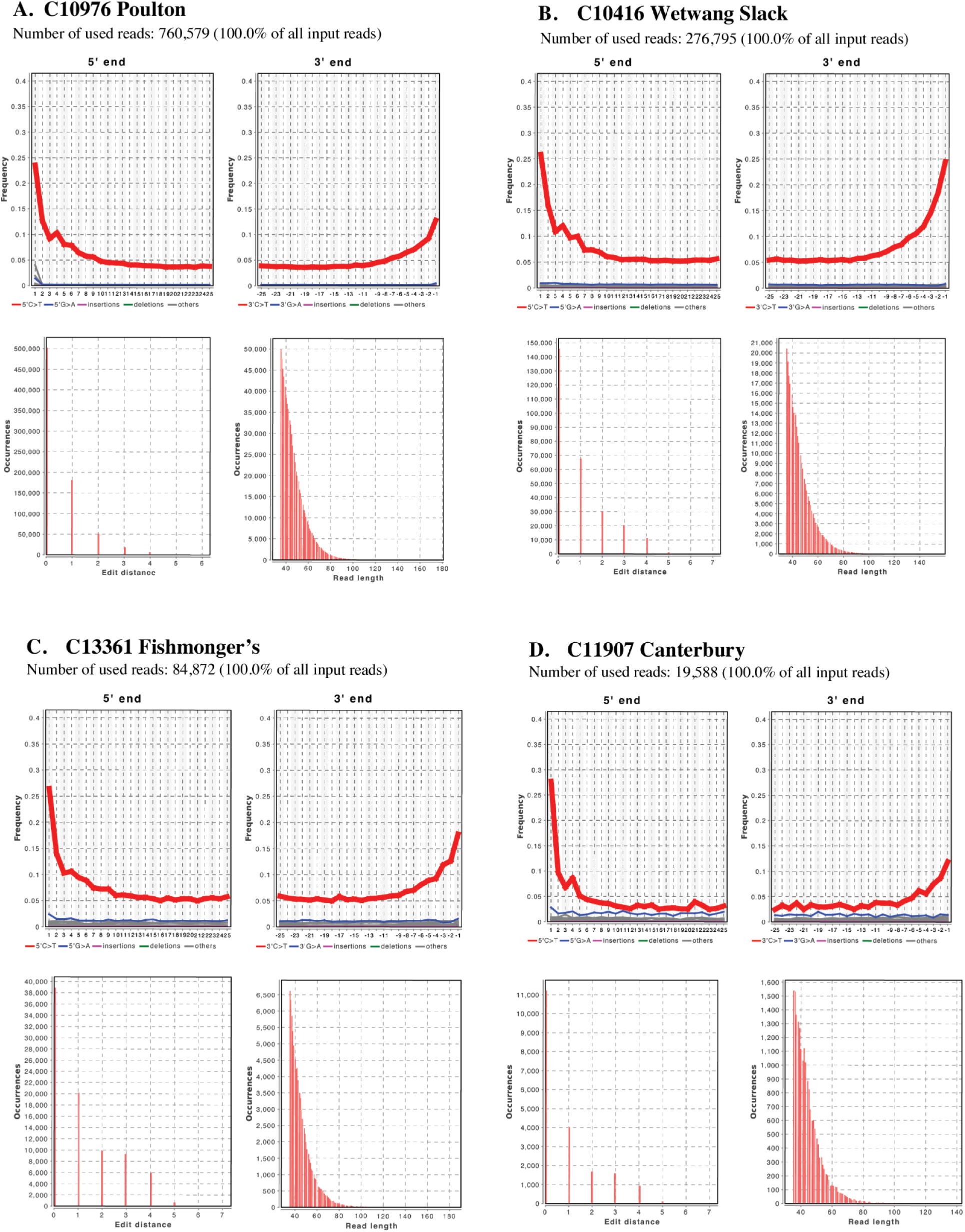
Ancient pathogen authentication of four ancient genomes from this study when aligned to *B. recurrentis* A1 reference. Edit distance, Damage and fragment length distribution for the concatenated genomes for each sample when aligned to *B. recurrentis* chromosome and plasmids (A1 reference genome) via DamageProfiler^53^. **a)** C10796 from Poulton **b)** C10416 from Wetwang Slack **c)** C13361 from Fishmonger’s **d)** C11907 from Canterbury.

**Figure S2.**
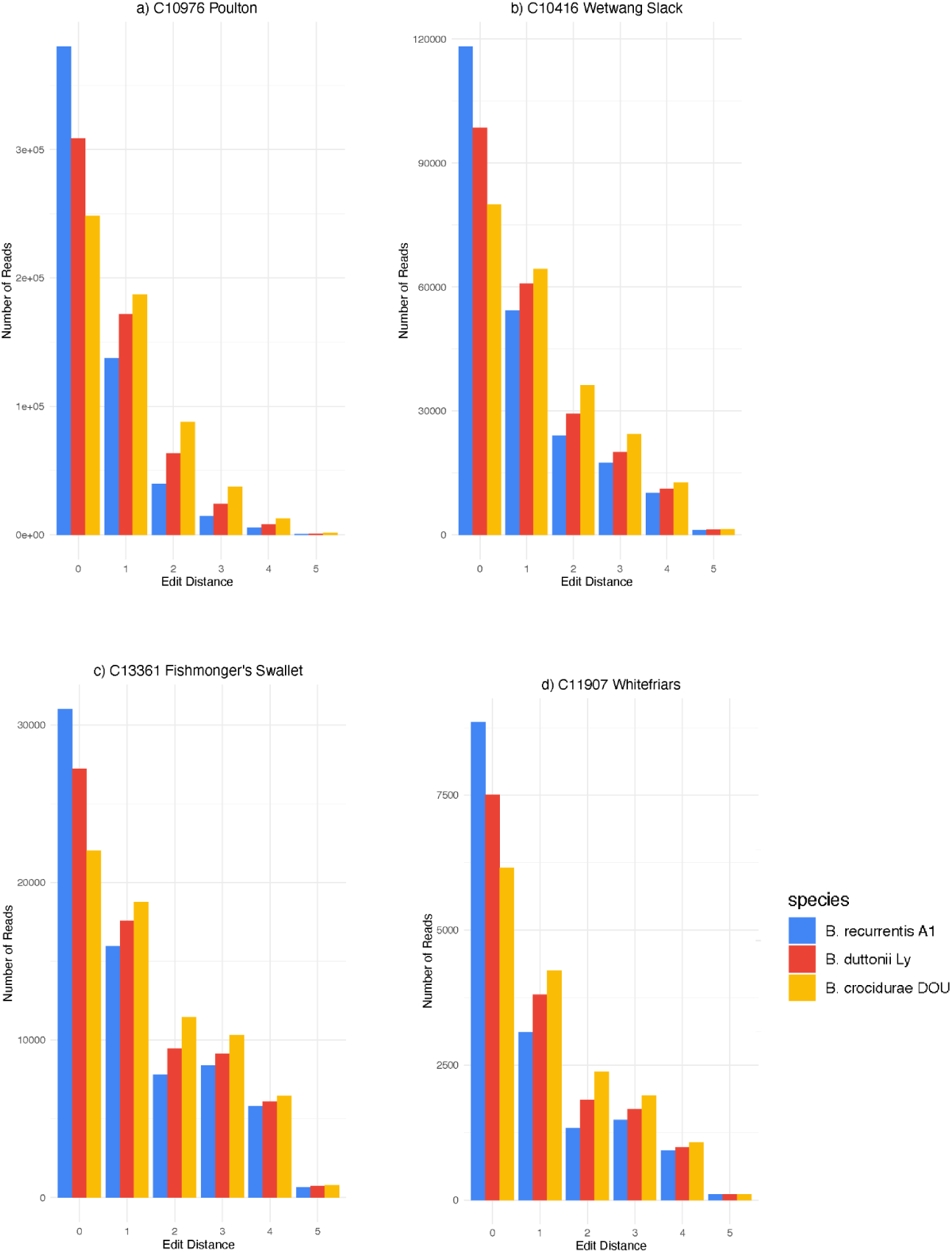
Edit distance of four ancient genomes from this study when aligned to *B. recurrentis* A1, *B. duttonii* Ly, and *B. crocidurae* DOU reference. Edit distance (x-axis) against number of reads (y-axis) of all newly considered ancient genomes from this study, when aligned to *B. recurrentis* A1 (blue)*, B. duttonii* Ly (red) and *B. crocidurae* DOU (yellow) reference genomes. **a)** C10796 from Poulton **b)** C10416 from Wetwang Slack **c)** C13361 from Fishmonger’s **d)** C11907 from Canterbury

**Figure S3.**
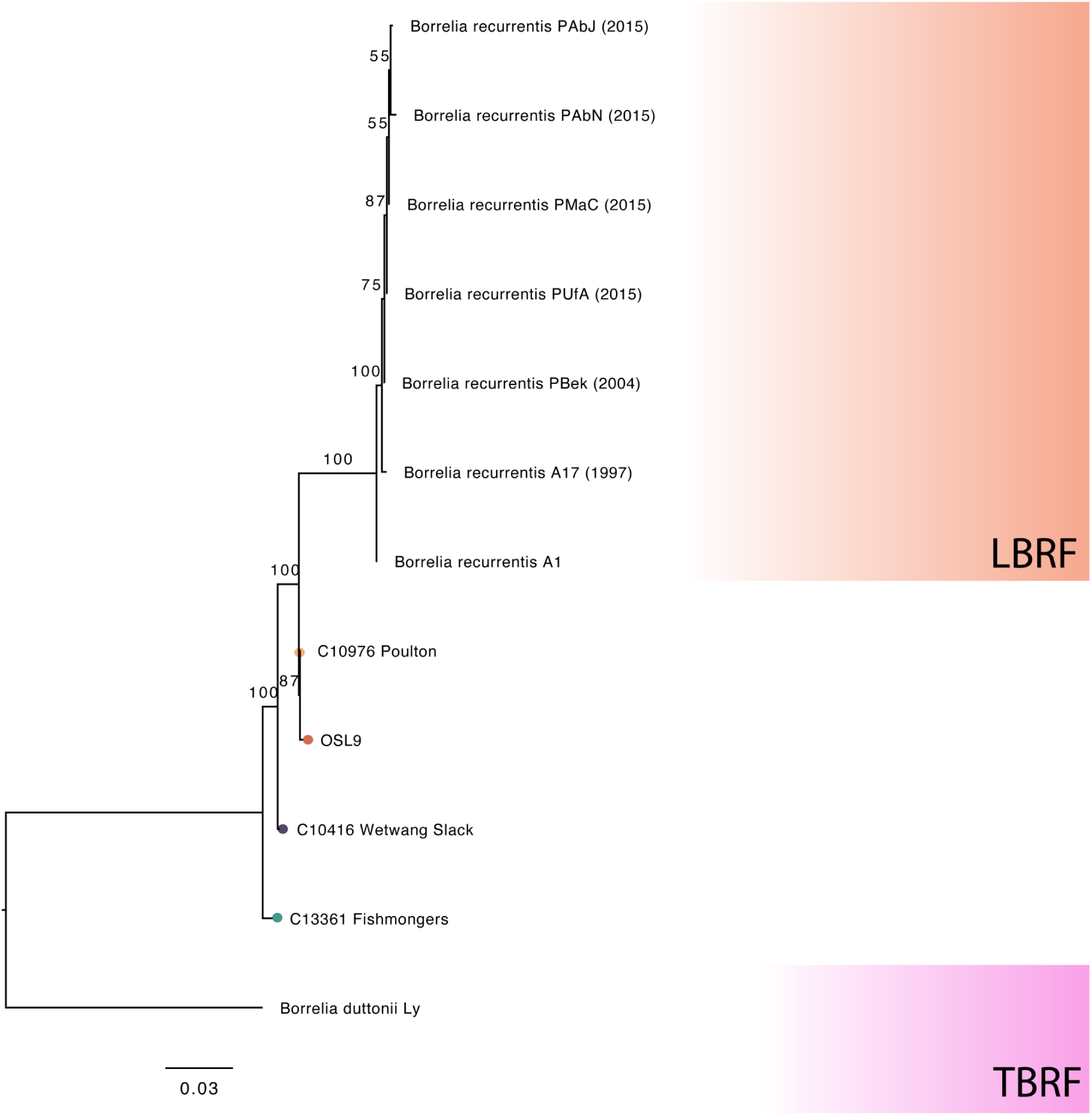
Maximum Likelihood SNP phylogeny of ancient and modern *B. recurrentis* genomes and *B. duttonii* outgroup when aligned to Panaroo core-genome. Maximum likelihood phylogenetic tree (GTR+F+ASC according to AIC) constructed on variability over the core genome SNP alignment of 2,006 sites filtering for 20% missing per site and per genome including transitions.

**Figure S4.**
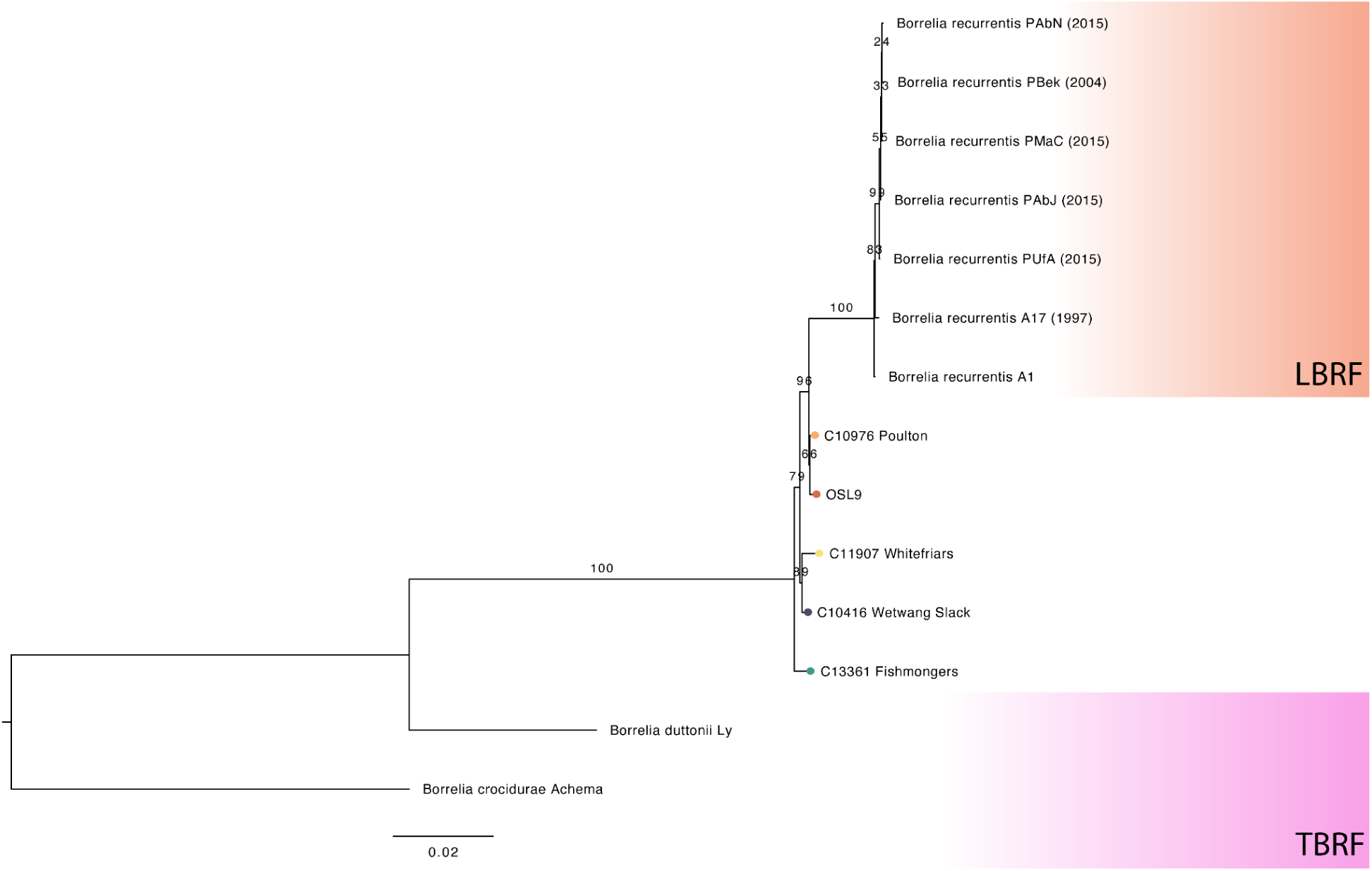
Maximum Likelihood SNP phylogeny of ancient and modern *B. recurrentis* genomes and *B. duttonii Ly* outgroup when aligned to *B. recurrentis* A1 reference genome with relaxed filtering thresholds. Maximum likelihood phylogenetic tree (K3Pu+F+ASC model implemented with rapid bootstrapping) constructed on variability over a reference based alignment to the *B. recurrentis* A1 reference genome (3,354 sites, 20% missingness per site, maximum missingess across genome 90%). All ancient genomes were filtered for a minimum base fold coverage of 3 except for C11907 Canterbury which was filtered for a minimum coverage of 2.

**Figure S5.**
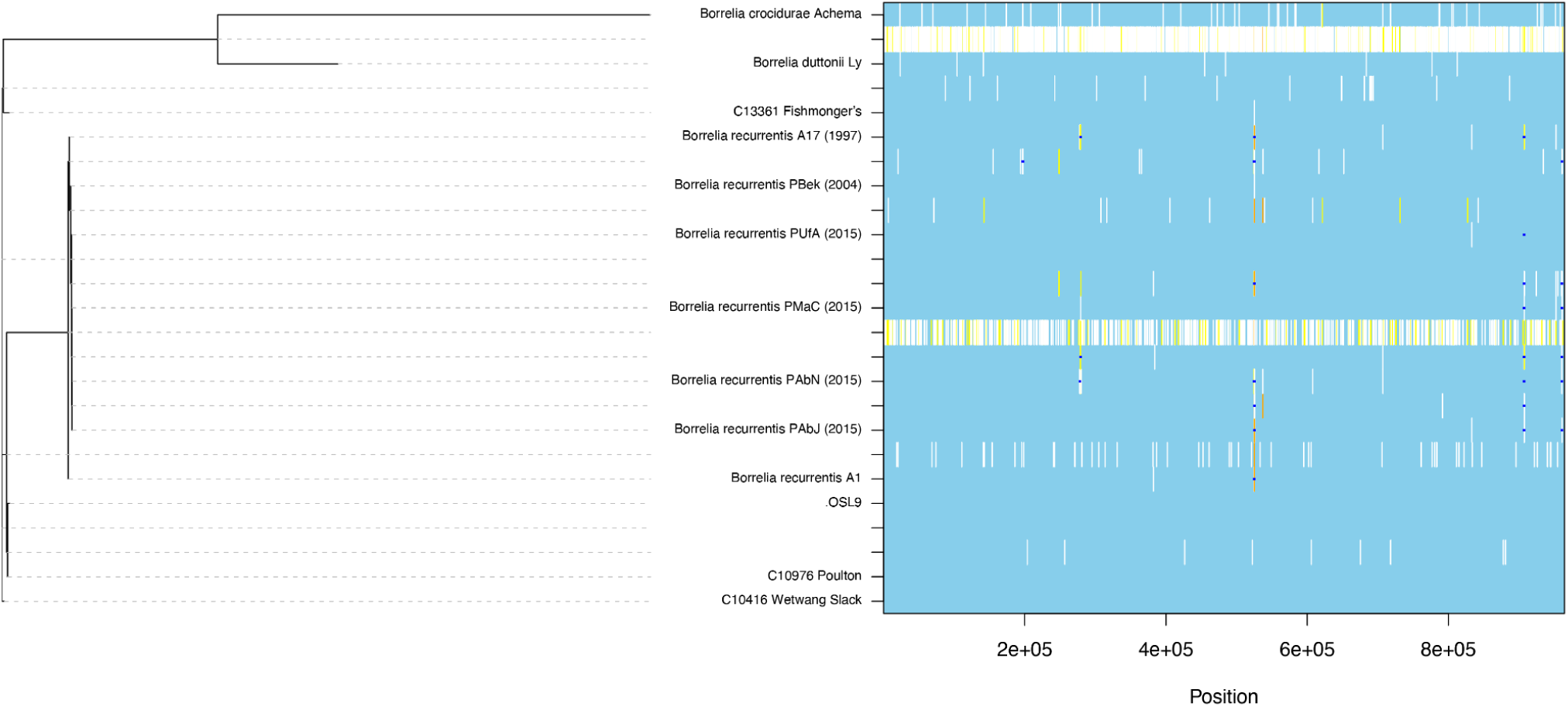
ClonalFrameML homoplasy recombination analysis of core genome. Reconstructed substitutions (white vertical bars) are shown for each branch of the maximum likelihood tree. Dark blue horizontal bars indicate recombination events detected by the analysis with yellow providing homoplasic positions.

**Figure S6.**
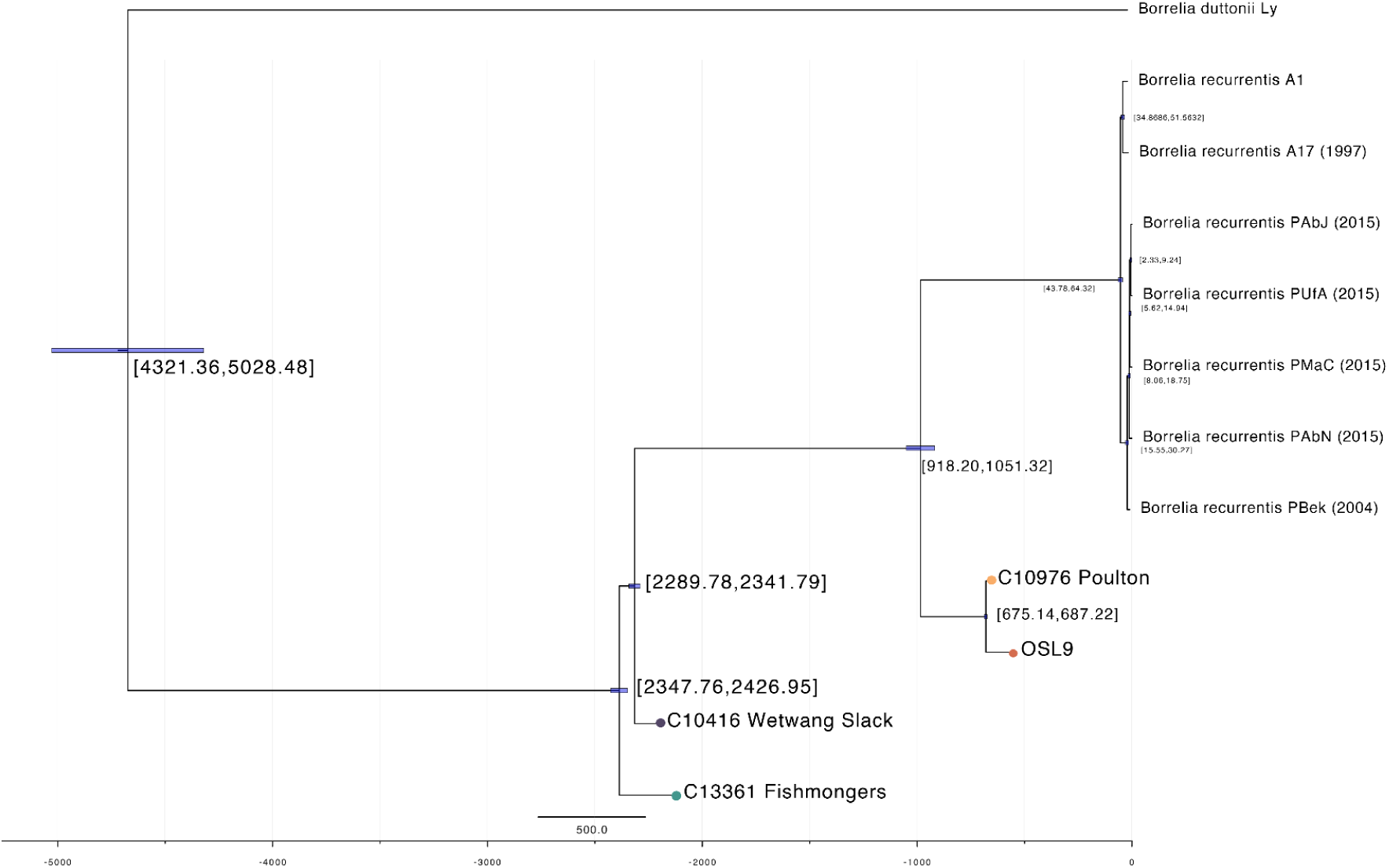
Beast2 tip-calibrated time tree when C13361 Fishmonger’s genome is included. Bayesian tip-calibrated maximum clade credibility time tree from Beast2, providing the best-supported model following path-sampling when C13361 Fishmonger’s is included in the alignment. Confidence intervals around nodes provided the 95% higher posterior density. Ancient samples are highlighted by coloured tips.

**Figure S7.**
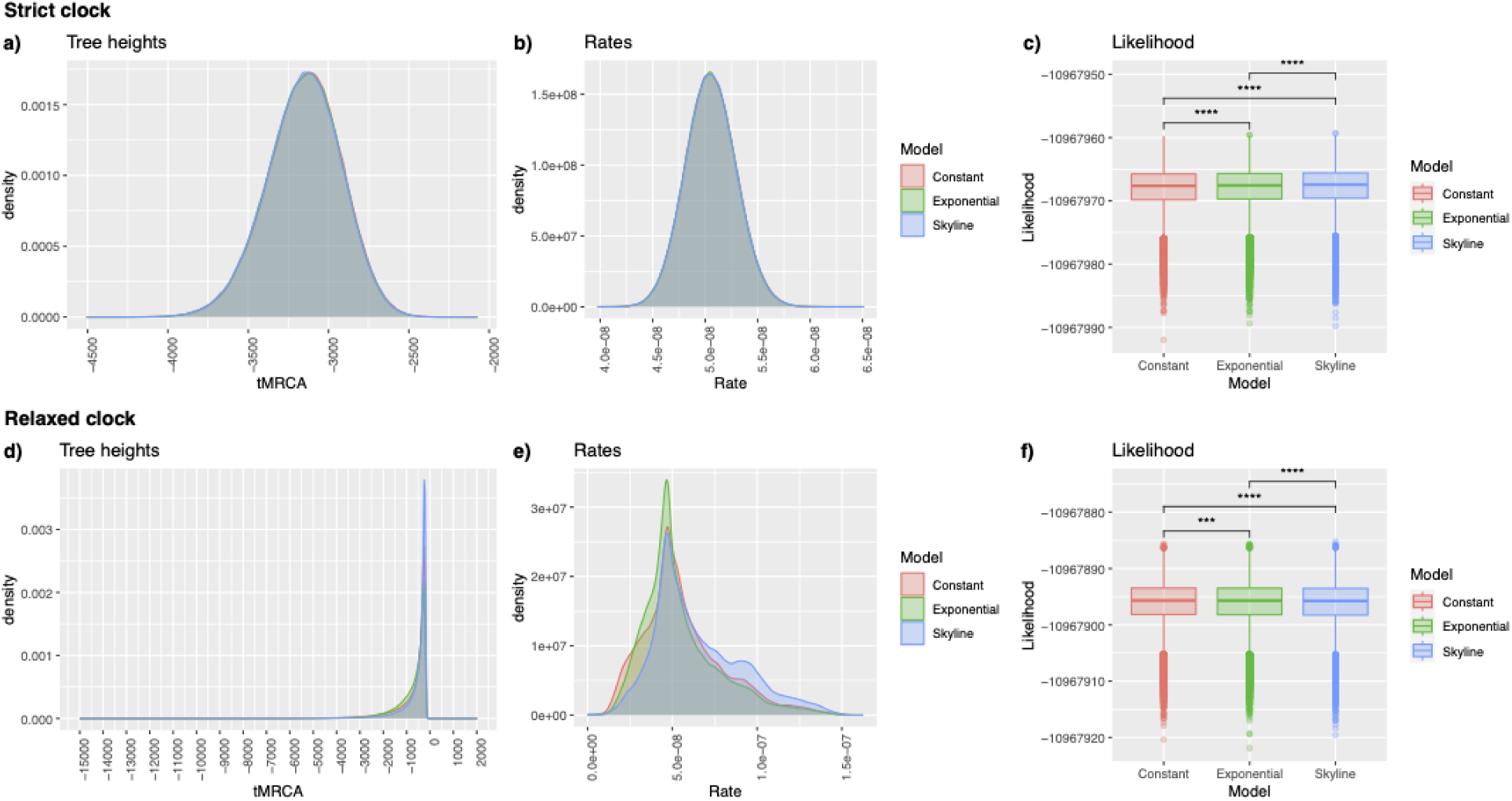
Posterior Distributions from Bayesian tip-dating analysis. Posterior distributions estimated under a strict clock (top row) and relaxed clock (bottom row) allowing three possible specifications of demographic models (red - coalescent constant, green - coalescent exponential, blue - coalescent skyline). Estimated tree heights are given in the left-hand panels (a,d), estimated clock rates are provided in the right-hand panels (b,e). Boxplots provide the posterior distribution of likelihood values under a strict (c) and relaxed (f) clock model.

**Figure S8.**
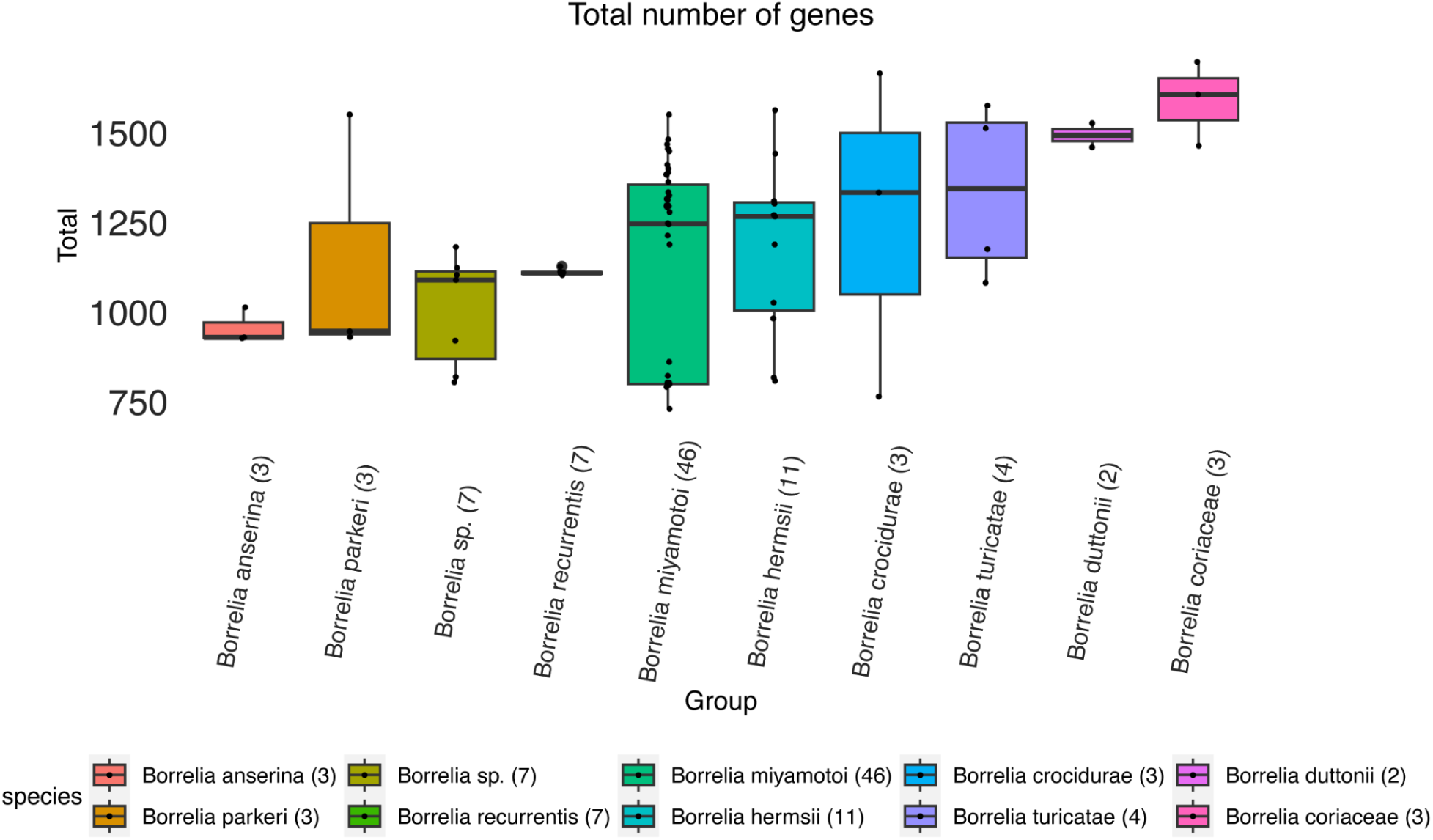
Per species distribution of the total number of genes identified across the *Borrelia-wide* pan-genome after filtering for truncated genes. The count of the number of genomes considered per species is provided in parentheses. Genes were filtered for truncated genes using the Panaroo *–filter* parameter. Species where only one isolate was available was excluded from this plot.

**Figure S9.**
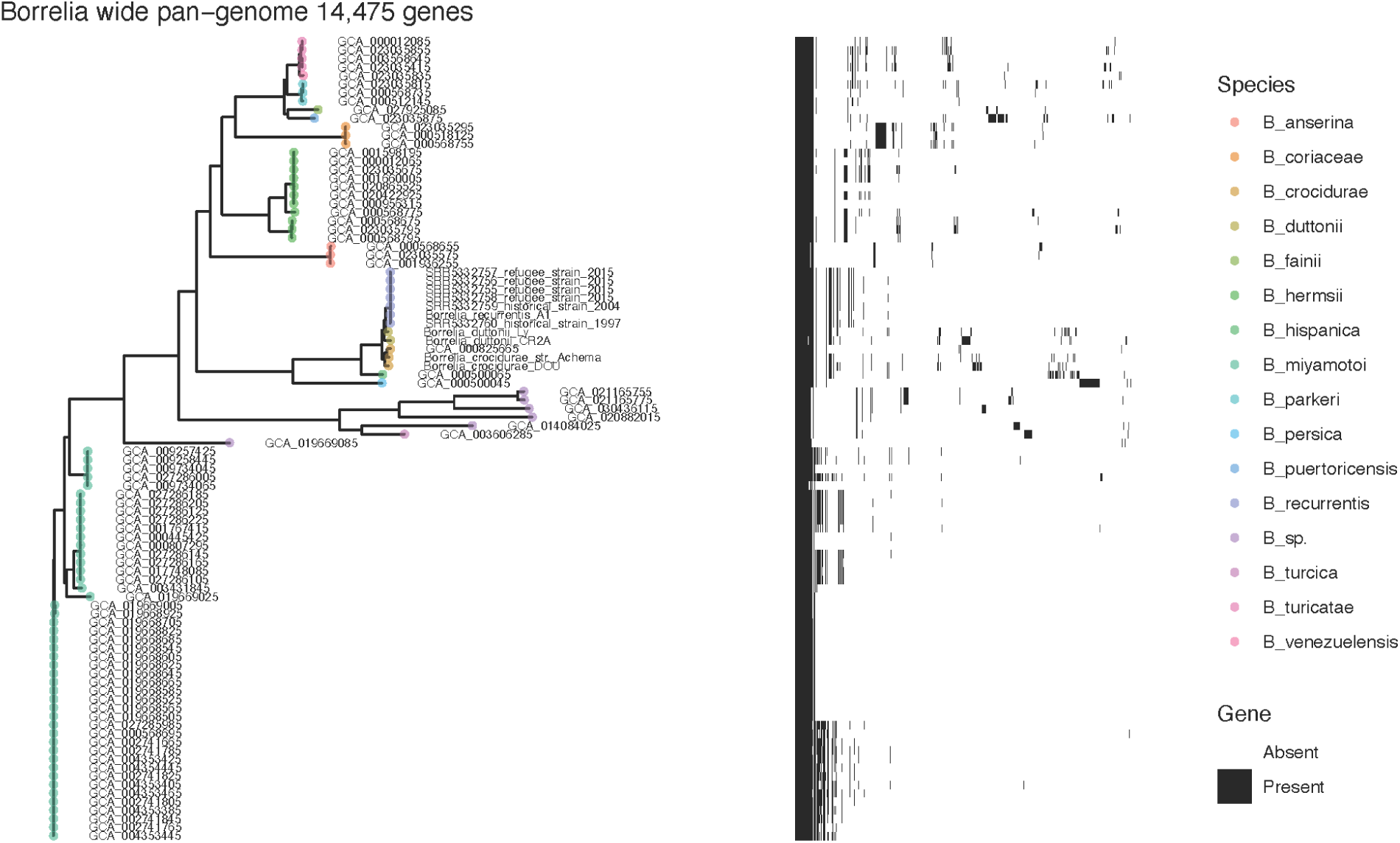
Pan-genome diversity across the *Borrelia* (RF) genus, comprising 14,475 genes. The phylogeny at right provides a neighbour-joining representation based on the gene presence and absence over the 14,475 pan-genome. Heatmap at right provides the presence (black) and absence (white) across the pan-genome, with each column providing an individual gene.

**Figure S10.**
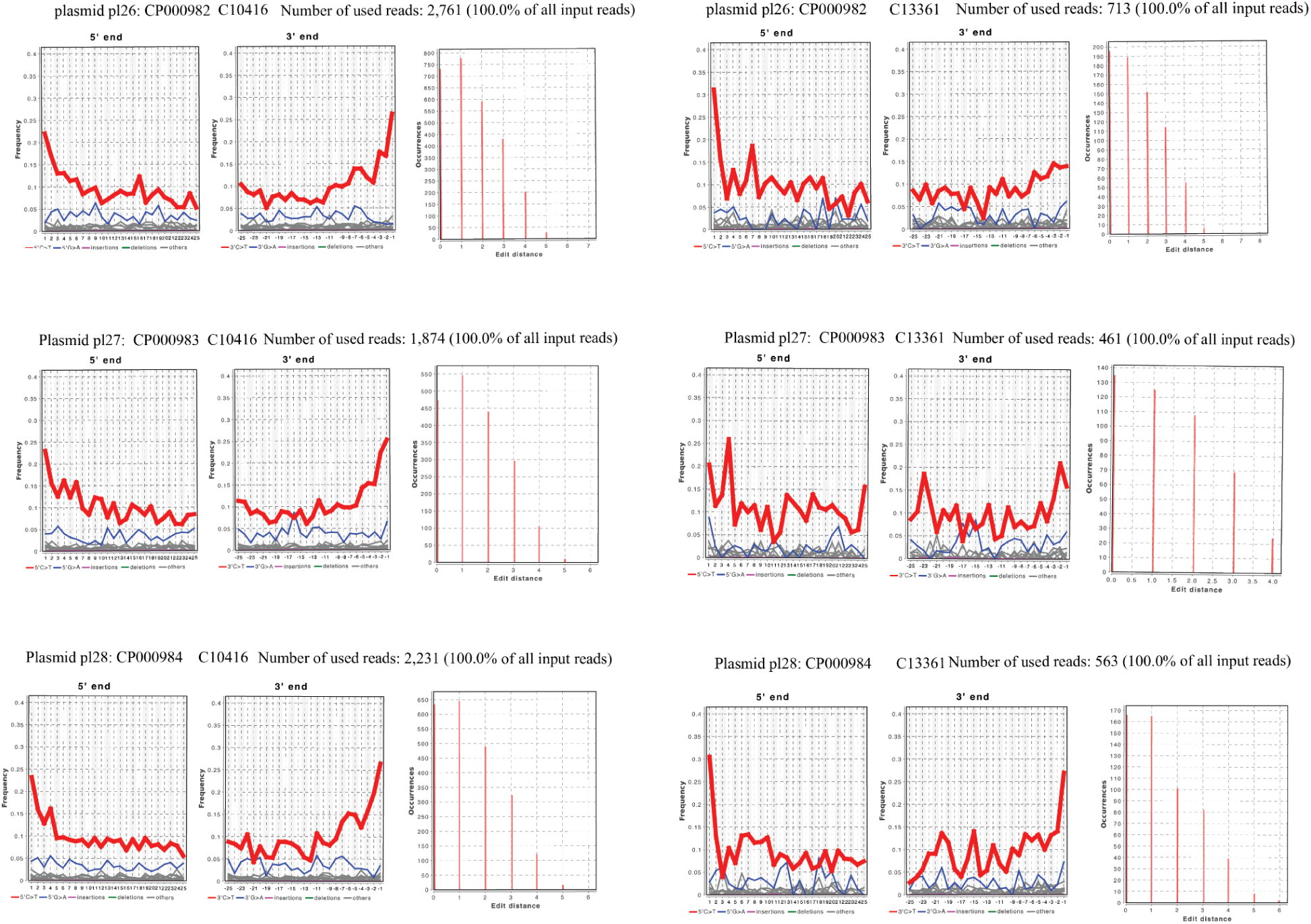
Plasmid authentication of *B. duttonii* plasmids pl26, pl27, pl28 in C10416 Wetwang Slack and C13361 Fishmonger’s. Damage (deamination) pattern and edit distance of C10416 Wetwang Slack and C13361 Fishmonger’s when aligned to *B. duttonii* plasmids pl26, pl27 and pl28 which are shown to be higher coverage than all other *B. recurrentis* genomes.

**Figure S11.**
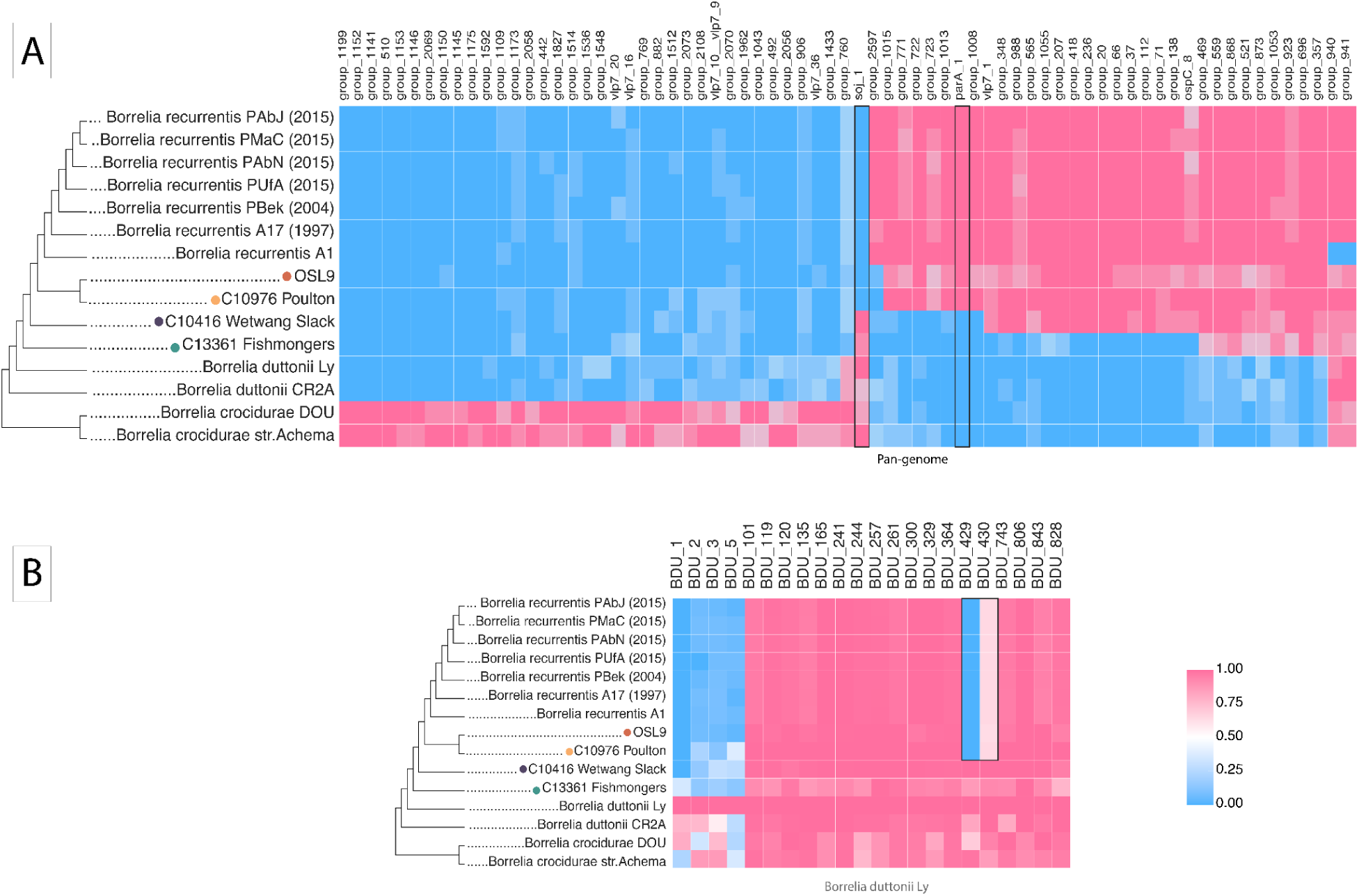
Presence/absence across pan-genome genes and gene clusters. Ancient and modern genomes were aligned to the pan-genome reference built on all modern *B. recurrentis*, *B. duttonii* and *B. Crocidurae* genomes (**Methods**). Normalised coverage across the gene and gene clusters was calculated using BEDtools v2.29.2. Genes that had a coverage between 0.3 and 0.7 were filtered out and genes that has a consensus coverage across the clade modern *Borrelia*, medieval clade was kept. Additionally, genes that are in the same state (either all absent or all present) in the *B. duttonii* Ly and ancient and modern *B. recurrentis* were also filtered out. This resulted in 71 genes out of the 3035 genes identified in the pan-genome output. Regions of interest highlighted in text are outlined with a black box. Cladogram provides the relationship between different genomes based on a SNP phylogeny

**Figure S12.**
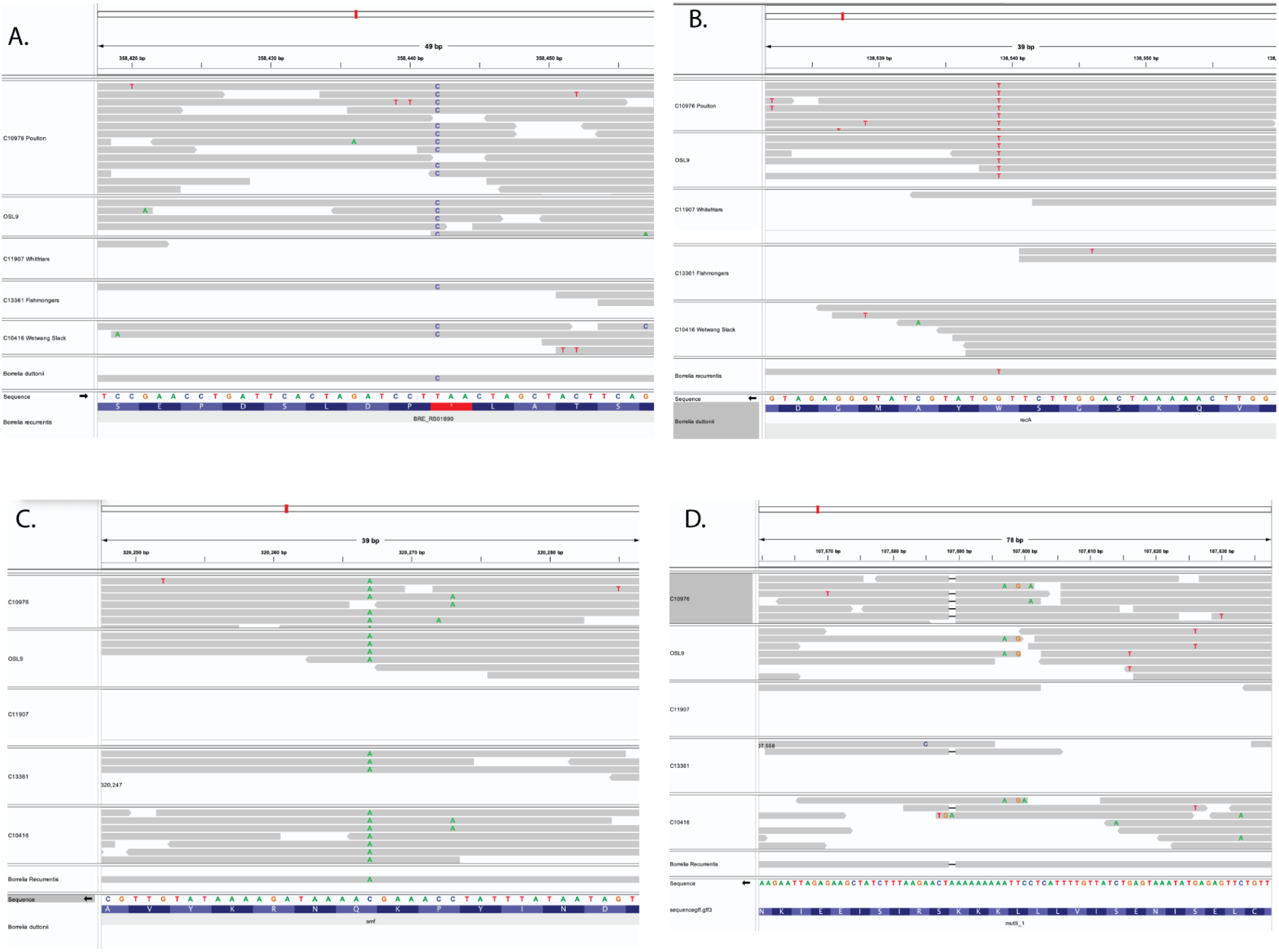
Integrated Genome Viewer of SNPs and INDELs. Previously identified SNPs and INDELS shown to be in different states in *B. recurrentis* A1 and *B. duttonii* Ly, visualised with IGV 2.17.4. **A)** Ancestral state of a polymorphism in the *oppA-1* gene in ancient samples from this study when aligned to *B. recurrentis* A1. The derived allele, which is present in *B. recurrentis,* results in an inframe stop codon. **B)** In-frame stop codon in the *recA* gene in *B. recurrentis* and medieval genomes, from a Tryptophan TGG (W) in *B. duttonii Ly* to a TAG stop codon (seen as a T on the forward strand), in the *B. recurrentis* A1 genome and the two medieval genomes included in this study. C10416 Wetwang Slack confidently shows the functional form of the gene, as does C11907 Canterbury but with the support of only 1 read. **C)** In-frame stop codon in the *smf* gene. IGV plot showing A on the forward strand in *B. recurrentis* A1 and all ancient genomes, resulting in a TAA in the reverse strand leading to an in-frame stop codon. The *B. duttonii* sequence is shown in the direction of protein synthesis. **D)** A frameshift mutation in the *mutS* gene, which is present in *B. recurrentis*, is represented by a deletion. Here we show the presence of reads aligning to the *mutS* gene with the same frameshift mutation present in *B. recurrentis A1*, also present in C10416 Wetwang Slack, C13361 Fishmonger’s, and C10976 Poulton. Due to the low number of reads aligning to this region and the potential misalignment of the reads, it is unclear as to whether the frameshift mutation is absent or present in the other ancient individuals, or whether their presence is an artefact caused by misalignment.

**Figure S13.**
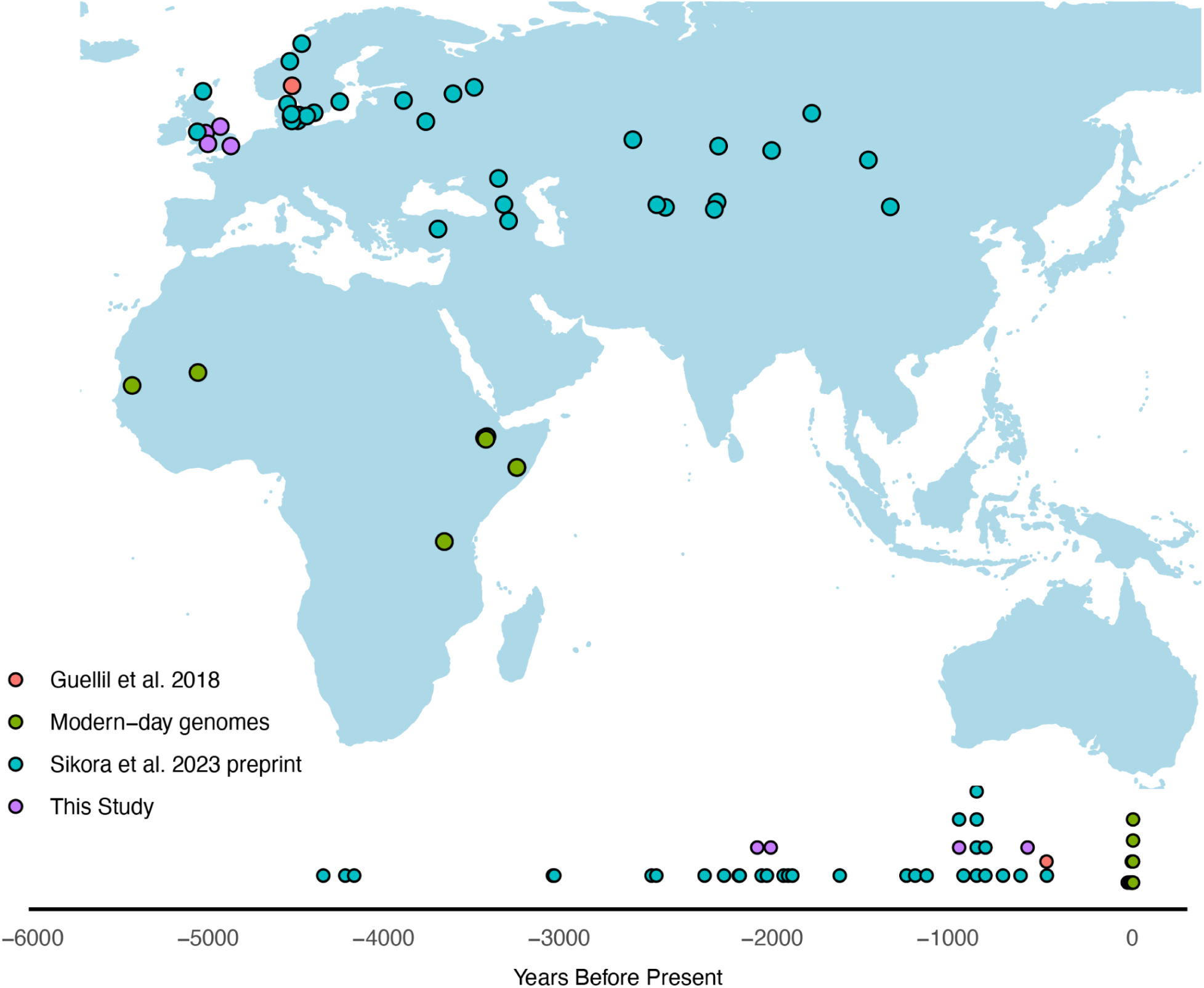
Map and timeline showing the distribution of ancient and modern *B. recurrenti*s genome through space and time. Map providing the geographic location and timeline of *B. recurrentis* observations to date. Note that observations from Sikora et al. 2023 preprint comprise hits assessed from partial recovery of sequencing reads as opposed to whole genome observations.

**Figure S14.**
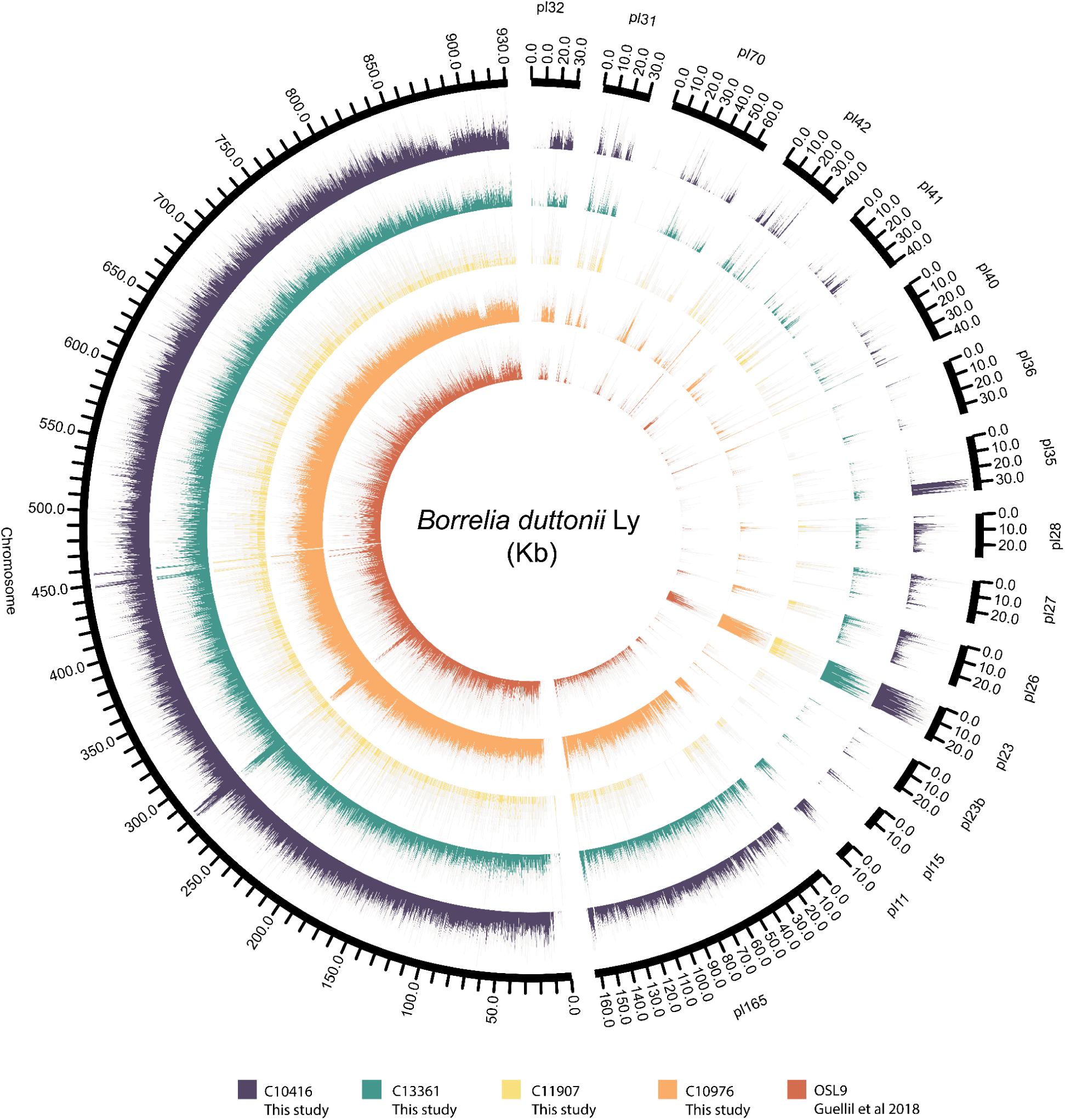
Circos plots providing the per sample coverage across the *B. duttonii Ly* reference genome. Circos plot providing the coverage of ancient genomes across the *B. duttonii* chromosome and plasmids when aligned to the *B. duttonii Ly* reference genome (GCF_000019685.1). A window size of 100bp for the chromosome and 10bp for the plasmids was used to provide the normalised coverage per window plotted.

**Figure S15.**
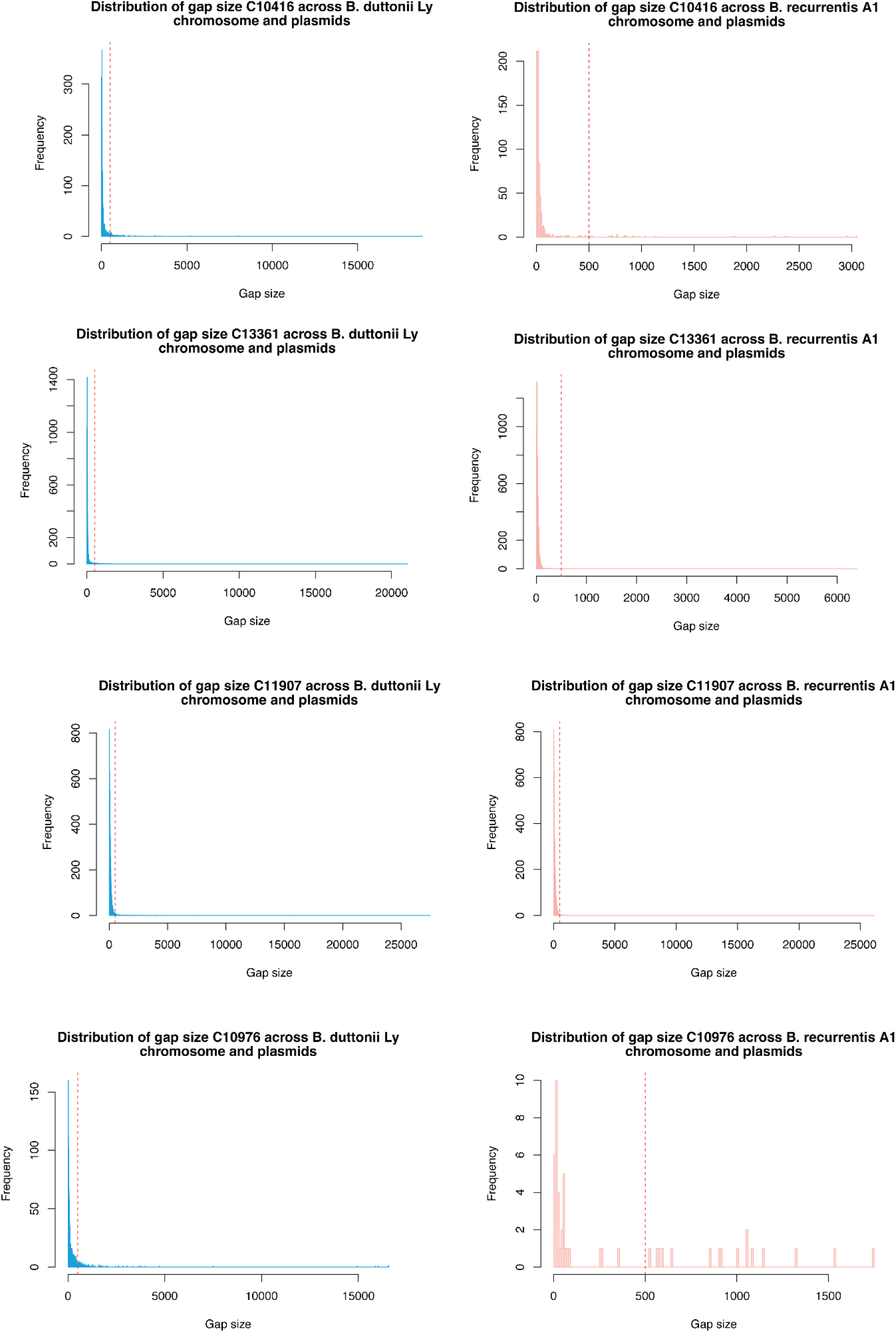
Gap size distribution plot to identify deletions over 500bp. Missingness distribution across ancient individuals from this study when aligned to the *B. duttonii* Ly chromosome (blue) and *B. recurrentis* A1 chromosome and plasmids (pink). Positions of missingness window above 500bp (red dashed line) are reported in **Table S8**.

## Supplementary Tables

Table S1 All modern and ancient genomes used in this analysis.

Table S2 Assessment of temporal signal following the BactDating root-to-tip evaluation procedure.

Table S3 BEAST2 posterior probability estimates following assessment of the results over three demographic models tested using both strict and relaxed clock priors.

Table S4 Number of SNPs identified as ’Ancestral’ and ’Derived’ in ancient genomes

Table S5 Coverage across plasmids when aligned to *B. duttonii* Ly reference genome

Table S6 Normalised coverage after filtering for the 71 genes post filtering showing temporal patterning across our dataset

Table S7 Identified gene ontology annotations in the 71 filtered genes and gene clusters from EggNoGG and interproScan

Table S8 Identified regions of missingness greater than 500bp when aligned to *B. recurrentis* A1 and B. duttonii Ly chromosome

